# CWF19L2 couples pre-mRNA alternative splicing with the maternal-to-zygotic transition to safeguard female fertility

**DOI:** 10.64898/2026.06.24.734384

**Authors:** Shiyu Wang, Tongtong Li, Yuling Cai, Kang Shangguan, Ziqi Wang, Chengqi Huang, Yajun Shi, Jing Xin, Qun Zhao, Honghui Zhang, Han Zhao, Yihong Guo, Hongbin Liu, Zi-Jiang Chen, Tao Huang

**Affiliations:** State Key Laboratory of Reproductive Medicine and Offspring Health, Center for Reproductive Medicine, Institute of Women, Children and Reproductive Health, Shandong University, 250012, China; Center for Reproductive Medicine, The First Affiliated Hospital of Zhengzhou University, Zhengzhou, 450052, China; National Research Center for Assisted Reproductive Technology and Reproductive Genetics, Shandong University, Jinan, Shandong, 250012, China; Key Laboratory of Reproductive Endocrinology (Shandong University), Ministry of Education, Jinan, Shandong, 250012, China; Shandong Technology Innovation Center for Reproductive Health, Jinan, Shandong, 250012, China; Shandong Provincial Clinical Research Center for Reproductive Health, Jinan, Shandong, 250012, China; Shandong Key Laboratory of Reproductive Research and Birth Defect Prevention, Jinan, Shandong, 250012, China; Research Unit of Gametogenesis and Health of ART-Offspring, Chinese Academy of Medical Sciences (No.2021RU001), Jinan, Shandong, 250012, China; Advanced Medical Research Institute, Cheeloo College of Medicine, Shandong University, Jinan, Shandong, 250012, China; Henan Key Laboratory of Reproduction and Genetics, The First Affiliated Hospital of Zhengzhou University, Zhengzhou, 450052, China; Center for Reproductive Medicine, Department of Reproductive Endocrinology, Zhejiang Provincial People’s Hospital (Affiliated People’s Hospital), Hangzhou Medical College, Hangzhou, Zhejiang, 310014, China; Shanghai Key Laboratory for Assisted Reproduction and Reproductive Genetics, Shanghai, China; Department of Reproductive Medicine, Ren Ji Hospital, Shanghai Jiao Tong University School of Medicine, Shanghai, China

**Author notes:** Correspondence (Y. Guo); (H. Liu); (Z.-J. Chen); (T. Huang). These authors contributed equally to this work (Shiyu Wang, Tongtong Li, Yuling Cai).

**Keywords:** CWF19L2, alternative splicing, female infertility, oocyte and early embryo competence defects (OECD), maternal-to-zygotic transition (MZT)

## Abstract

The maternal-to-zygotic transition (MZT) requires precise spatiotemporal execution of pre-mRNA alternative splicing (AS), yet the core splicing machinery driving this developmental reprogramming remains incompletely understood. Here, we identify the CWF19-like protein 2 (CWF19L2) as an indispensable pre-mRNA AS regulator that safeguards against oocyte and early embryo competence defects (OECD). While murine germline-specific depletion of *Cwf19l2* spares morphological folliculogenesis, oocyte maturation, or fertilization, it induces complete female sterility characterized by profound developmental arrest at the 2-cell stage. Mechanistically, maternal deficiency of CWF19L2 localized to nuclear speckles disrupts transcription-splicing-translation coupling, collapsing AS homeostasis during maternal reserves and zygotic genome activation. We further demonstrate that CWF19L2 orchestrates the pre-mRNA splicing network through directly binding to target transcripts and indirectly modulating via interacting with the core spliceosomal factor PRPF8. Importantly, exogenous *Cwf19l2* mRNA partially rescues the embryonic arrest. Together, our findings establish CWF19L2 as an indispensable AS engine during the MZT, providing a mechanistic foundation for OECD and a novel molecular etiology for female infertility.

## Introduction

Successful female reproduction hinges upon a highly orchestrated sequence of events: oocyte maturation, fertilization, and early embryonic development^[1]^. During oocyte maturation, the oocyte synthesizes and accumulates essential maternal transcripts and proteins, and then undergoes extensive transcriptional silencing^[2, 3]^. Following fertilization, the newly formed zygote still remains temporarily transcriptionally silent^[4]^ and relies entirely on stored maternal factors to trigger the maternal to zygotic transition (MZT)^[5]^. MZT represents a gradual shift of early embryonic development control from the maternal genome to the zygotic genome^[6]^, characterized by maternal substance clearance and zygotic genome activation (ZGA)^[7]^. ZGA unfolds in two waves: the initial minor ZGA represents the earliest transcriptional event, followed by the major ZGA that triggers large-scale transcriptional activity^[6, 8]^. In mice, minor ZGA typically occurs at the late 1-cell (zygotic) stage, while major ZGA takes place at the late 2-cell stage; in humans, these two events occur at the 4-cell stage and 8-cell stage, respectively^[9]^. Following ZGA, the embryo progressively establishes autonomous regulation over its developmental trajectory^[10]^. This highly coordinated transition serves as the fundamental prerequisite for subsequent embryogenesis and successful gestation.

Any aberration in oocytes and embryos may give rise to developmental defects or miscarriage^[11]^. Clinically, oocyte and early embryo competence defects (OECD) represent a severe class of female infertility due to compromised oocyte quality, frequently culminating in recurrent in vitro fertilization (IVF) failures with no transferable embryos across multiple cycles^[12]^. OECD exhibits profound phenotypic heterogeneity, including empty follicle, oocyte maturation arrest, fertilization failure, zygote arrest, early embryonic arrest (EEA), and mixed phenotypes, with EEA emerging as the most prevalent. However, currently established genetic determinants collectively explain only approximately 20% of cases. The etiology for OECD patients remains elusive^[13, 14]^, creating a critical bottleneck in both clinical diagnosis and our fundamental understanding of oogenesis and embryogenesis. Therefore, an in-depth dissection of the mechanisms underlying governing the oocyte-to-embryo transition is essential for advancing and safeguarding human reproductive health.

Alternative splicing (AS) represents a crucial post-transcriptional RNA processing regulatory layer that exponentially expands transcriptomic and proteomic diversity by generating multiple mRNA isoforms from a single gene^[15, 16]^. The precise execution of AS is executed by the spliceosome, a highly dynamic macromolecular complex comprising over 170 proteins that orchestrate a sequential cycle of assembly, activation, catalysis, and disassembly to link exons and remove introns^[17–19]^, which exists in ten different complexes throughout the reaction^[20, 21]^. During MZT, mRNA is subjected to extensive AS dynamics, and this process is referred to as zygotic splicing activation (ZSA), a crucial event coupled temporally with ZGA in both humans and mice^[22]^. Transcriptomic profiling reveals that the 2-cell embryos exhibit the highest frequency of mRNA alternative splicing events (ASEs) across preimplantation development^[23, 24]^, underscoring the absolute necessity of precise isoform expression for successful embryogenesis.

As interest in the modulation of AS during MZT continues to rise, numerous genetic studies have revealed the important roles of splicing factors in regulating oogenesis and embryogenesis, such as BCAS2, DDX5, hnRNPM, ESRP1, PCBP1, RTCB, SRSF1, SRSF2 (SC35), SRSF3, SF3B1, and YTHDC1^[22, 25–37]^. These findings further illustrate the significant impact of AS on OECD. For instance, the ablation of BCAS2, a core component of the spliceosomal PRP19 complex, disrupts spliceosome assembly and consistently precipitates oocyte and embryo developmental arrest and apoptosis by mis regulating the AS of key developmental genes (e.g., *Dazl*, *Pabpc1l*, and *Nobox*)^[25–28]^. And hnRNPM can cooperate with BCAS2, modulating its binding to pre-mRNA loci to precisely regulate AS^[30]^. Similarly, SRSF family proteins act as essential gatekeepers of reproductive success: SRSF1 governs primordial follicle formation via regulating the AS of POI-related genes^[27]^; SRSF2 reactivation in early embryos is pivotal for ZSA^[22, 38]^; and SRSF3 deficiency uniformly induces early embryonic arrest by inducing aberrant AS^[34]^. However, the overarching regulatory architecture and the precise molecular mechanisms governing AS dynamics during the MZT remain largely elusive.

The evolutionarily conserved splicing factor CWF19L2 (CWF19-like cell cycle control factor 2) has previously been implicated male fertility. Its deficiency impairs spermatogenesis, a defect driven by not only directly binding to and regulating the AS of genes related to essential genes, but also amplifying splicing regulation effect via RBFOX1^[39]^. In addition, structural discoveries revealed by cryo-electron microscopy show that in the late stages of splicing, CWF19L2 is recruited to interact with PRPF8 to form the intron lariat spliceosome (ILS) complex^[40]^. Beyond reproduction, chromosomal CWF19L2 deletion is identified in breast carcinomas^[41]^, and CWF19L2 has positive associations with the risk of coronary artery calcification^[42]^. While the structural and part of the mechanism of CWF19L2 are known, its physiological role and mechanism in the female reproductive system remain further investigation.

In this study, we demonstrate that CWF19L2 is a critical orchestrator of AS during oogenesis and embryogenesis. Spatiotemporal analysis reveals that CWF19L2 exhibited stage-specific nuclear localization throughout folliculogenesis, vanished during meiotic resumption, and returned at the 2-cell stage, aligning with transcription and splicing activation. The female germline conditional *Cwf19l2* knockout mice are absolutely sterile due to embryonic arrest at the 2-cell stage, despite exhibiting normal oocyte morphological maturation, ovulation, and fertilization. At the molecular level, CWF19L2 deletion disrupts the transcription-splicing-translation coupling, inducing abnormalities in genes related to splicing, cell cycle and others, which resulted in poor oocyte and early embryo quality. Mechanistically, CWF19L2 directly regulates the AS networks of genes involved in AS regulation and ZGA, and physically interacts with PRPF8, acting as a key co-factor, to indirectly amplify splicing efficiency. Collectively, our findings confirm the indispensable function of CWF19L2 in regulating AS during MZT, thereby safeguarding female reproductive capacity.

## Results

### Spatiotemporal dynamics of CWF19L2 within the nuclear speckles of oocytes and early embryos

The evolutionarily conserved protein CWF19L2 has been previously documented to exhibit high expression within the reproductive system and to exert a pivotal role in male spermatogenesis^[39]^. To delineate its function in females, we first profiled CWF19L2 expression in multiple organs of mice via immunoblotting and qPCR. The results revealed that both the protein and mRNA levels of CWF19L2 were conspicuously elevated in the ovary (Figure 1A and S1A). Concomitantly, ovarian CWF19L2 expression displayed a fertility-correlated dynamic pattern: it escalated from pubescence (3-week-old) to reproductive maturity (6-week and 2-month-old), declined progressively alongside the age-related diminution of fertility (6 to 8-month-old) (Figure 1B and S1A).

**Figure 1.**
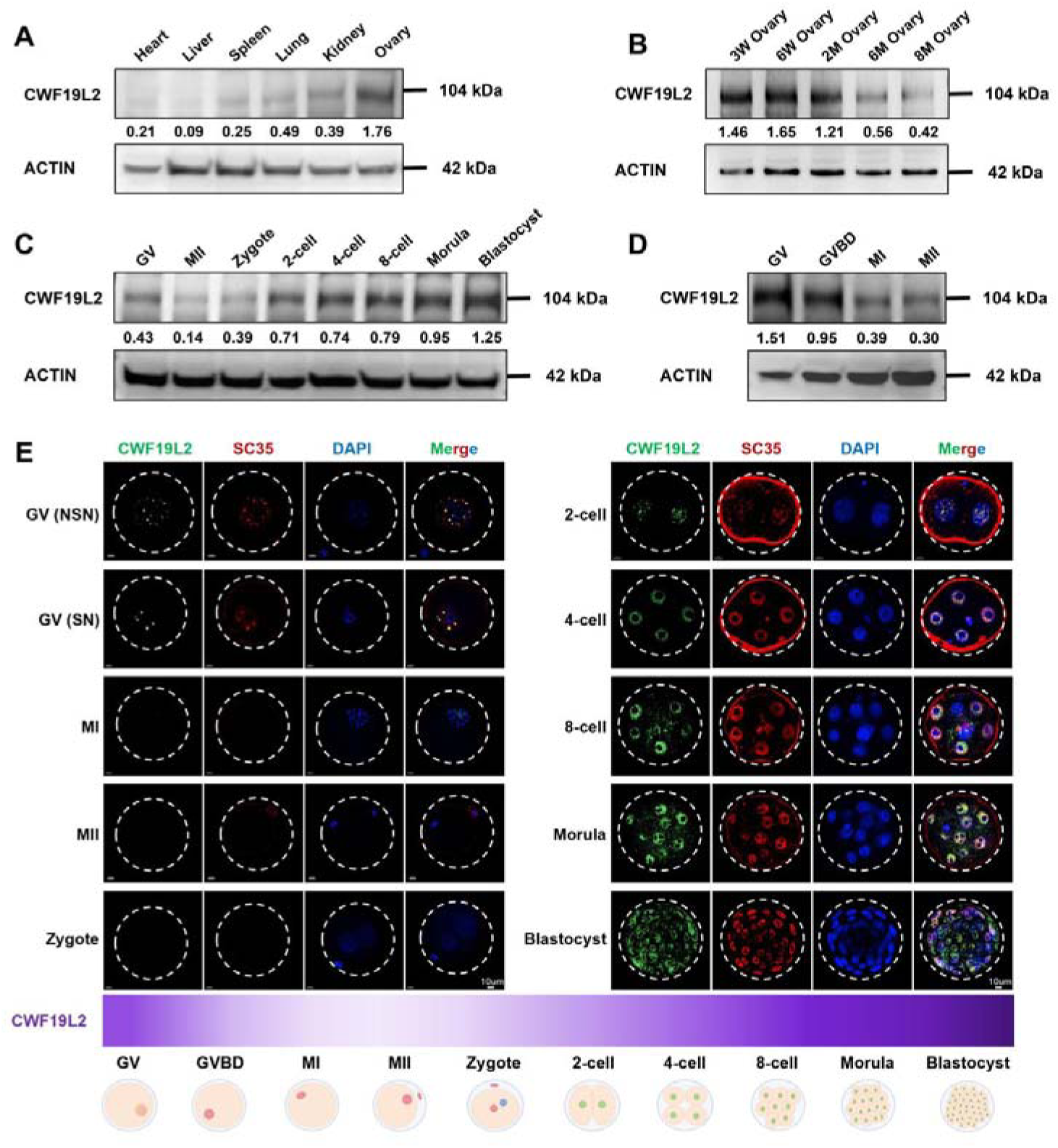
Spatiotemporal expression profiling of CWF19L2 during oogenesis and early embryogenesis **A.** Immunoblotting analysis of CWF19L2 in multiple organs of adult wild-type (WT) mice. ACTIN served as the loading control. Relative quantification is shown in the middle panel. **B.** Immunoblotting analysis of CWF19L2 at different stages of ovaries in WT mice. ACTIN served as the loading control. Relative quantification is shown in the middle panel. **C-D.** Immunoblotting analysis of CWF19L2 at different stages of oocytes and embryos in WT mice. ACTIN served as the loading control. Relative quantification is shown in the middle panel. **E.** Co-immunofluorescence staining of CWF19L2 (green) and the nuclear speckles marker SC35 (red) in different stages of oocytes and embryos of WT mice. DNA was stained with DAPI. The expression pattern of CWF19L2 is shown below. Scale bars = 10 µm.

Next, we characterized the spatiotemporal distribution of CWF19L2 during oocyte maturation, fertilization, and early embryonic development by immunoblotting, qPCR, and immunofluorescence. Notably, as transcription is progressively silenced in germinal vesicle (GV) oocytes, CWF19L2 expression gradually declined; conversely, following fertilization, it was progressively re-enhanced as embryos developed (Figure 1C-1D and S1B). Co-immunofluorescence with SC35 (the nuclear speckles marker) revealed that CWF19L2 punctually localized within the nucleus and shared nearly identical expression and localization with SC35 (Figure 1E). Specifically, CWF19L2 forms numerous diminutive speckles in non-surrounded nucleolus (NSN) oocytes with active transcription and splicing, and these speckles transitioned to larger, sparser aggregates in quiescent surrounded nucleolus (SN) oocytes. Following a transient disappearance during meiosis and fertilization, prominent CWF19L2 speckles re-emerged at the 2-cell embryo and intensified during subsequent development, precisely mirroring the onset of ZGA and ZSA (Figure 1E and S1C-L). This dynamic expression pattern closely matches the paradigm of maternal factors, which control early events of embryonic development.

Collectively, these observations underscore that CWF19L2 is widely expressed in oocytes and embryos with active splicing, implicating it in the post-transcriptional control indispensable for female fertility.

### CWF19L2 interplays with PRPF8 through the CWFJ domain

During pre-mRNA splicing, the spliceosome undergoes a dynamic P-to-ILS complex transition to release ligated exons^[43, 44]^. Structurally, PRPF8 serves as the central scaffold anchoring splicing factors during transition, sequentially dissociating SLU7 and recruiting CWF19L2 and PRPF43^[43, 45–47]^. Consistent with this structural hierarchy, in silico analysis using the STRING database predicted that CWF19L2 may interact with PRPF8, PRPF43, SLU7 and SC35 (Figure 2A). And we confirmed these interactions in HEK293T cells via co-immunoprecipitation (Co-IP) followed by immunoblotting (Figure 2B). Reciprocal Co-IP further confirmed that PRPF8 and PRPF43 associated with CWF19L2, with PRPF8 showing the higher enrichment efficiency (Figure 2C, S2A-C). Using Proximity Ligation Assay (PLA), we also unequivocally confirmed the direct incorporation between CWF19L2 and PRPF8 in HEK293T cells (Figure 2D). To dissect the mechanistic basis of their interaction, we ectopically co-expressed CWF19L2 and PRPF8 in HEK293T cells and mapped the critical binding domains via reciprocal Co-IP assays. Truncation mapping revealed that the 100-811aa region (containing the N-domain), the 812-1752aa region (containing the Linker and endonuclease domain), and the 1766-2020aa region (containing the RNaseH-like domain) of PRPF8 are indispensable for its association with CWF19L2 (Figure 2E). Conversely, CWF19L2 interacts with PRPF8 through its 657-779aa region (containing the CWF-J-1 domain) and 788-882aa region (containing the CWF-J-2 domain) (Figure 2F).

**Figure 2.**
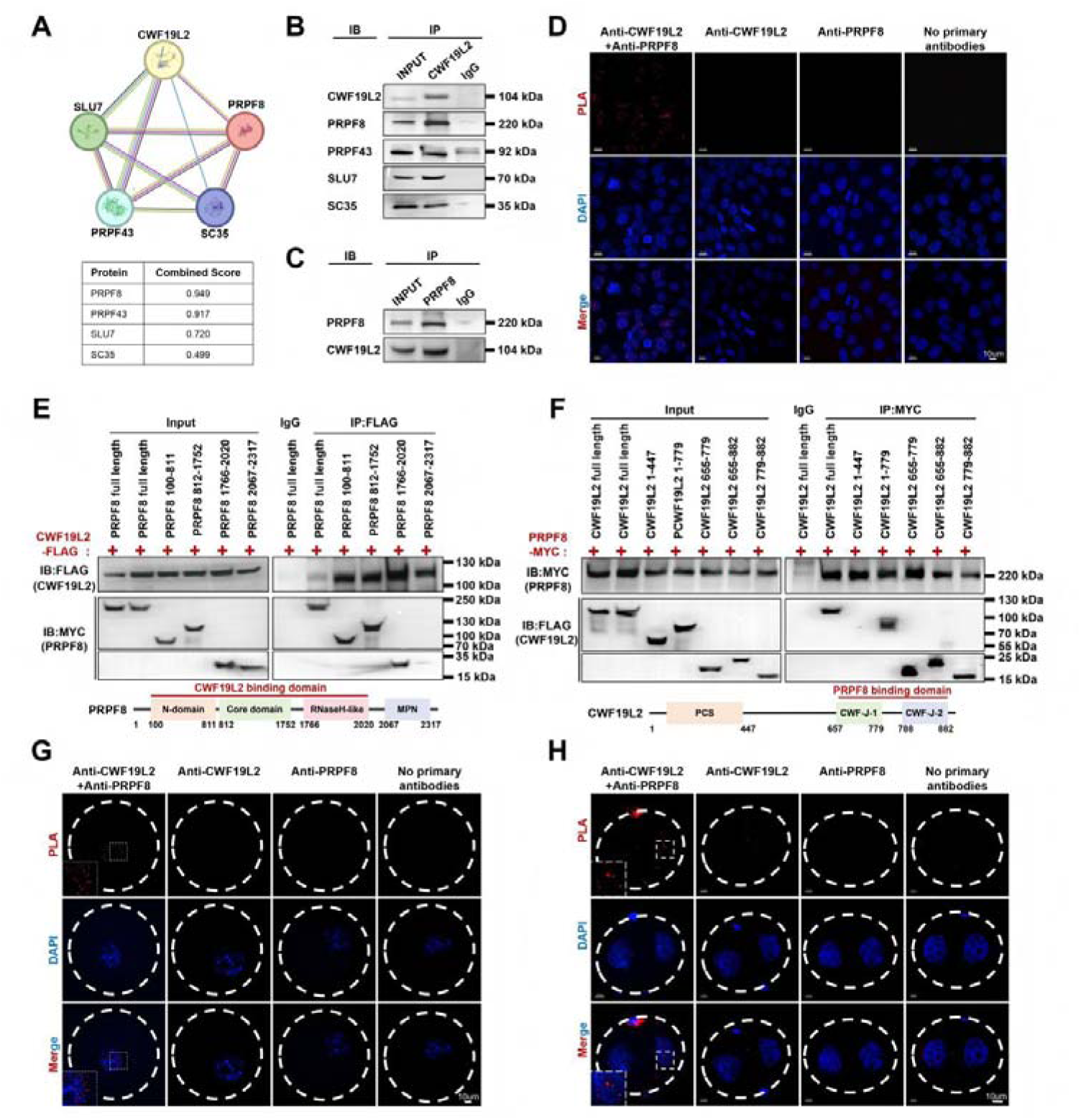
CWF19L2 physically interacts with PRPF8 through the CWFJ domain **A.** STRING database analysis shows the predicted interaction network of CWF19L2 with 4 mRNA-splicing-related proteins. **B.** Validation of the interactions between CWF19L2 and PRPF8, PRPF43, SLU7, and SC35 in HEK293T cells by co-immunoprecipitation (Co-IP) assays. IgG was used as the negative control. **C.** Co-IP assays of the interaction between PRPF8 and CWF19L2 in HEK293T cells. IgG was used as the negative control. **D.** Proximity ligation assay (PLA) of the interaction between CWF19L2 and PRPF8 in HEK293T cells. Anti-CWF19L2, anti-PRPF8, and no-primary antibodies were used as the negative control. DNA was stained with DAPI. Scale bars = 10 µm **E.** Reciprocal Co-IP assays of interaction domains between CWF19L2 and PRPF8. HEK293T cells were co-transfected with CWF19L2-FLAG and the indicated fragments of PRPF8-MYC, immunoprecipitated with anti-FLAG antibody, and immunoblotted with FLAG and MYC antibodies. **F.** Reciprocal Co-IP assays of interaction domains between CWF19L2 and PRPF8. HEK293T cells were co-transfected with PRPF8-MYC and the indicated fragments of CWF19L2-FLAG, immunoprecipitated with anti-MYC antibody, and immunoblotted with MYC and FLAG antibodies. **G-H.** PLA of the interaction between CWF19L2 and PRPF8 in GV oocytes (G) and 2-cell embryos (H) of WT mice. Anti-CWF19L2, anti-PRPF8, and no-primary antibodies were used as the negative control. DNA was stained with DAPI. Scale bars = 10 µm

To contextualize these findings during maternal-to-zygotic transition, we performed co-immunofluorescence to interrogate the subcellular colocalization of CWF19L2 or SC35(the nuclear speckles marker) with SLU7, PRPF8 and PRPF43 in GV oocytes and 2-cell embryos. The results revealed their prominent colocalization within nuclear speckles where splicing occured (Figure S2D-G). Crucially, PLA once again confirmed the specific interaction between CWF19L2 and PRPF8 in GV oocytes and 2-cell embryos (Figure 2G-H). Analogously, we previously confirmed CWF19L2 interacted with PRPF8 and PRPF43 in the murine testes. Together, these observations supported the hypothesis that CWF19L2 may exert pre-mRNA splicing regulatory effects on oocytes and embryos through specific domain-dependent collaboration with some key splicing-related proteins, like PRPF8.

### CWF19L2 is indispensable for female fertility yet dispensable for morphological folliculogenesis in mice

To delineate the functional role of CWF19L2 in oogenesis, we generated mice harboring a floxed *Cwf19l2* allele, wherein exon 6 was flanked by loxP sites. By crossing *Cwf19l2*-floxed mice with *Stra8-GFPCre* transgenic mice (Figure 3A), we established germline-specific conditional *Cwf19l2* knockout mice (*Cwf19l2^flox/flox^; Stra8-GFPCre*, hereafter designated as “*Cwf19l2*-SKO” in the text or “SKO” in the figures). Notably, expression of *Stra8* in female germ cells initiates at embryonic day 13.0 (E13.0), peaks at E14.5, and reverts to normal levels by E16.5^[48, 49]^, which permits early and specific genetic excision. Genotypic validation was performed by PCR of tail biopsy DNA at postnatal day 5 (PD5) (Figure S3A). Subsequent quantitative qPCR, immunoblotting, and immunofluorescence analyses of oocytes consistently demonstrated the efficient depletion of both *Cwf19l2* mRNA and CWF19L2 protein in *Cwf19l2*-SKO mice relative to littermate controls (*Cwf19l2*^flox/+^ or *Cwf19l2*^flox/fox,^ hereafter referred as “control”) (Figure 3B-D).

**Figure 3.**
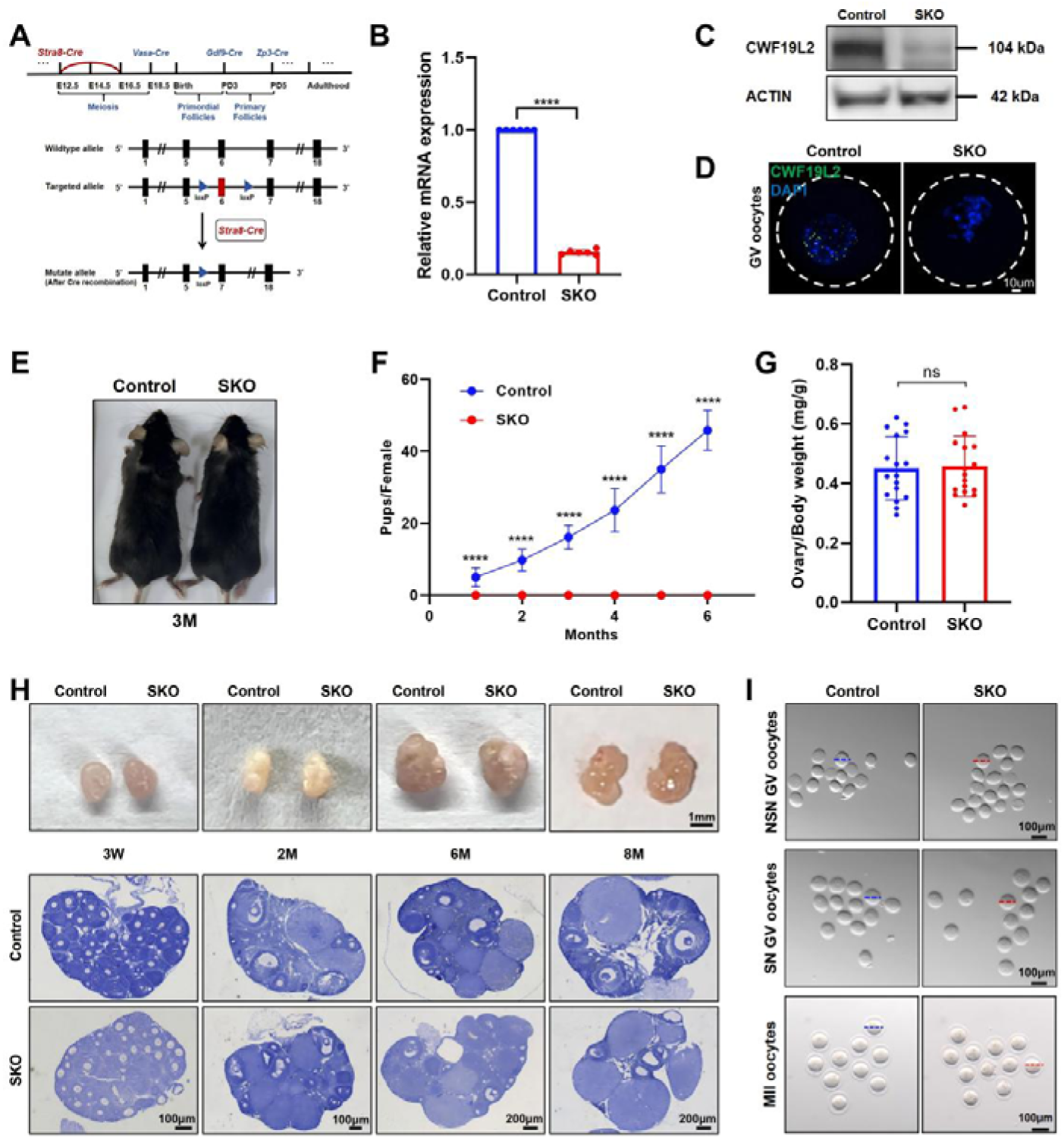
CWF19L2 is essential for female fertility but dispensable for morphological oocyte development **A.** The schematic showing the insertion of loxp sites to flank exon 6 of the mouse Cwf19l2 gene and deletion of Cwf19l2 in germ cells using Stra8GFP-Cre. **B.** QPCR analysis of knockout efficiency of Cwf19l2 in oocytes from Cwf19l2-SKO and control mice. Data are presented as mean ± SD, n = 6, ****:P < 0.0001. **C.** Immunoblotting analysis of the knockout efficiency of CWF19L2 in oocytes from Cwf19l2-SKO and control mice. ACTIN served as the loading control. **D.** Immunofluorescence analysis of the knockout efficiency of CWF19L2 (green) in oocytes from Cwf19l2-SKO and control mice. scale bars = 10 µm **E.** Morphology of adult Cwf19l2-SKO and control female mice. **F.** Cumulative number of pups per mouse during 6-month fertility tests of adult Cwf19l2-SKO and control mice. Data are presented as mean ± SD, n=3, ****:P < 0.0001. **G.** The ratio of ovary weight to body weight in adult Cwf19l2-SKO and control mice. Data are presented as the mean ± SD, n = 20, ns: not significant. **H.** Gross morphology (upper panel, scale bars = 1 mm) and hematoxylin staining (lower panel, scale bars = 100 or 200 µm) of ovaries from 3-week, 2-month, 6-month, and 8-month Cwf19l2-SKO and control mice. **I.** Representative images of NSN GV, SN GV, and MII oocytes obtained from Cwf19l2-SKO and control mice. Scale bars = 100 µm. The dotted lines represent diameters.

Although *Cwf19l2*-SKO female mice exhibited normal viability and appearance (Figure 3E), six-month fertility tests revealed they were completely sterile (Figure 3F). Intriguingly, this profound reproductive failure was uncoupled from macroscopic organ defects, as the ovary weight, body weight, and ovary-to-body weight ratio of *Cwf19l2*-SKO mice were indistinguishable from those of controls (Figure S3B-C and 3G). To investigate the causes of infertility due to CWF19L2 deficiency, we systematically analyzed ovarian and follicular development in *Cwf19l2*-SKO and control mice across distinct reproductive stages: 3 weeks (prepubescence), 2 months (early adulthood), 6 months (mid-reproductive age), and 8 months (late reproductive age). Comprehensive morphometric and histological examinations confirmed that mutant ovarian dimensions and the quantities of primordial, primary, secondary, antral follicles, and corpus luteum were completely comparable to controls across all age cohorts (Figure 3H and S3D). Given the above, CWF19L2 serves as an indispensable factor for female fertility in mice, whereas it is unnecessary for the morphological establishment of follicles and ovaries.

### Depletion of CWF19L2 precipitates embryonic development arrest at the 2-cell stage

Subsequently, we extended our investigations to interrogate whether oocyte development and quantity were perturbed in *Cwf19l2*-SKO mice. To this end, we implemented a stage-specific oocyte collection strategy: NSN oocytes were isolated from 2-week-old mice, SN oocytes from 3-week-old mice, and MII oocytes from 2-month-old mice following administration of pregnant mare serum gonadotropin (PMSG) and human chorionic gonadotropin (hCG). Detailed microscopic assessments revealed no discernible disparities in oocyte phenotypic appearance between *Cwf19l2*-SKO and control mice (Figure 3I). Furthermore, the number and diameter of NSN, SN and MII oocytes retrieved from *Cwf19l2*-SKO mice were comparable to those of control groups (Figure S3E). Together, these data indicate that CWF19L2 depletion does not appreciably compromise the quantity or morphological maturation of oocytes.

Given the normal gametogenic progression, we sought to unravel the underlying cause of sterility in *Cwf19l2*-SKO mice by assessing developmental competence post-fertilization. We performed IVF using MII oocytes from *Cwf19l2*-SKO and control mice paired with wild-type sperm, generating maternal *Cwf19l2* mutant (*Cwf19l2*^♀-/♂+^) and control embryos (*Cwf19l2*^♀+/♂+^). Continuously monitoring embryonic development in culture revealed that while oocytes derived from *Cwf19l2*-SKO mice were successfully fertilized and progressed to the 2-cell stage at rates equivalent to controls, nearly all failed to advance beyond this stage (Figure 4A-C). By 96 hours after fertilization, maternal mutant embryos arrested at the 2-cell stage exhibited overt degenerative features, including increasing granular material and worsening cytoplasmic blebbing and fragmentation (Figure 4A). These comprehensive observations underscore an indispensable role for CWF19L2 in early embryonic development, implicating its critical involvement in driving MZT.

**Figure 4.**
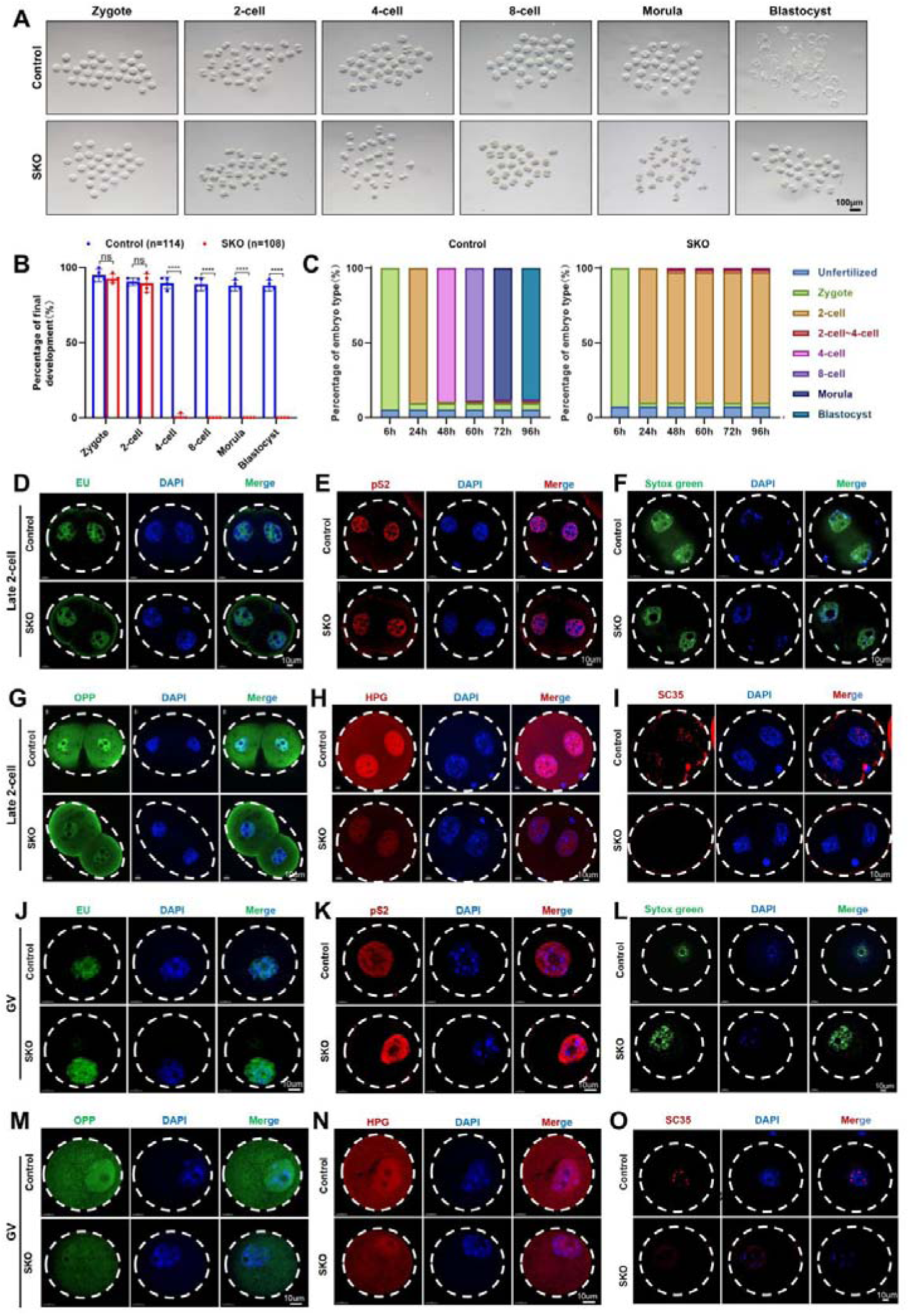
Depletion of CWF19L2 precipitates embryonic development arrest at the 2-cell stage **A.** Representative images of pre-implantation embryos at different stages derived from adult Cwf19l2-SKO and control mice. Scale bars = 100 µm. **B.** Quantification of pre-implantation embryos at different stages derived from adult Cwf19l2-SKO (n=108) and control mice (n=114) at 96h after fertilization. Data are presented as the mean ± SD. ****:P < 0.0001, ns: not significant. **C.** The percentage composition of different embryo types in adult Cwf19l2-SKO and control mice at 6, 24, 48, 60, 72 and 96 h after fertilization. **D.** Immunofluorescent staining of EU showing RNA transcription in late 2-cell embryos from Cwf19l2-SKO and control mice. Scale bars = 10 µm. **E.** Immunofluorescent staining of pS2 showing RNA transcription in late 2-cell embryos from Cwf19l2-SKO and control mice. Scale bars = 10 µm. **F.** Distribution of DNA and RNA in late 2-cell embryos from Cwf19l2-SKO and control mice. DAPI stains DNA only. Sytox green stains both DNA and RNA. **G.** Immunofluorescent staining of OPP showing protein synthesis in late 2-cell embryos from Cwf19l2-SKO and control mice. Scale bars = 10 µm. **H.** Immunofluorescent staining of HPG showing protein synthesis in late 2-cell embryos from Cwf19l2-SKO and control mice. Scale bars = 10 µm. **I.** Co-immunofluorescence staining of CWF19L2 (green) with nuclear speckles marker SC35 (red) in late 2-cell embryos from Cwf19l2-SKO and control mice. DNA was stained with DAPI. Scale bars = 10 µm. **G.** Immunofluorescent staining of EU showing RNA transcription in GV oocytes from Cwf19l2-SKO and control mice. Scale bars = 10 µm. **K.** Immunofluorescent staining of pS2 showing RNA transcription in GV oocytes from Cwf19l2-SKO and control mice. Scale bars = 10 µm. **L.** Distribution of DNA and RNA in GV oocytes from Cwf19l2-SKO and control mice. DAPI stains DNA only. Sytox green stains both DNA and RNA. **M.** Immunofluorescent staining of OPP showing protein synthesis in GV oocytes from Cwf19l2-SKO and control mice. Scale bars = 10 µm. **N.** Immunofluorescent staining of HPG showing protein synthesis in GV oocytes from Cwf19l2-SKO and control mice. Scale bars = 10 µm. **O.** Co-immunofluorescence staining of CWF19L2 (green) with nuclear speckles marker SC35 (red) in GV oocytes from Cwf19l2-SKO and control mice. DNA was stained with DAPI. Scale bars = 10 µm.

### CWF19L2 deficiency triggers embryonic arrest through disrupting transcription-splicing-translation coupling

During MZT, the embryo needs to initiate the synthesis of nascent RNA and proteins to sustain subsequent development^[5]^. Abnormalities in ZGA represent the most prevalent causes of embryonic developmental arrest at the 2-cell stage^[6, 9]^. We therefore assessed the expression of multiple ZGA-related genes in late 2-cell embryos, and found substantial downregulation of these hallmark genes in CWF19L2-deficient embryos (Figure S4A), which may serve as a critical driver of EEA.

To dissect the underlying mechanism of embryonic arrest in *Cwf19l2*-SKO mice, we first evaluated global transcriptional activity in 2-cell embryos using two complementary approaches: 5-ethynyluridine (EU) incorporation assays (de novo RNA synthesis) and immunofluorescence of RNA polymerase II phosphorylated at serine-2 (pS2, the active transcription marker). These analyses revealed a modest reduction in transcriptional activity in *Cwf19l2*-SKO embryos relative to controls (Figure 4D-E and S5A-B). We next assessed global RNA accumulation by co-staining 2-cell embryos with Sytox Green (labeling both DNA and RNA) and DAPI (just labeling DNA), enabling simultaneous quantification of total nucleic acid content and DNA content. While the spatial distributions of Sytox Green and DAPI overlapped extensively in both *Cwf19l2*-SKO and control embryos (Figure 4F and S5C), quantitative analysis revealed stark differences. In controls, the average fluorescence intensity (AFI) of Sytox Green was significantly higher than that of DAPI, reflecting massive RNA accumulation consistent with robust ZGA-driven transcription. Whereas in *Cwf19l2*-SKO embryos, AFI of Sytox Green was only marginally elevated relative to DAPI (Figure S5D), resulting in a drastically depressed Sytox Green-to-DAPI fluorescence intensity ratio (Figure S5E), which demonstrated that *Cwf19l2*-SKO embryos fail to properly accumulate nascent RNA.

Next, we assessed global translational capacity using O-propargyl-puromycin (OPP) and L-homopropargylglycine (HPG) labeling nascent protein (Figure 4G-H). Strikingly, the relative OPP and HPG signal intensities in *Cwf19l2*-SKO.2-cell embryos were only 15-25% of those in control embryos (Figure S5F-G), indicating a near-complete collapse of translational activity.

Given that splicing serves as a critical nexus linking transcription and translation, we interrogated whether splicing was perturbed in *Cwf19l2*-SKO embryos. Immunofluorescence of SC35 yielded undetectable signals in *Cwf19l2*-SKO embryos (Figure 4I), suggesting that splicing initiation was also abrogated. Together, these results indicated that transcription was not fully reactivated in *Cwf19l2*-SKO 2-cell embryos, with downstream defects in splicing and translation, which collectively led to the failure of ZGA and ZSA, and ultimately resulted in embryonic arrest.

### CWF19L2 ablation uncouples transcription from splicing and translation during oogenesis

Inadequate maternal substance synthesis in oocytes constitutes another fundamental driver of EEA^[6, 13]^. Corroborating this, we detected dysregulated expression of multiple core maternal factors in GV oocytes (Figure S4B). We therefore conducted similar experiments as above in the GV oocytes (a critical stage for maternal material accumulation). The relative intensities of EU and PS2 in the *Cwf19l2*-SKO GV oocytes were higher than control counterparts (Figure 4J-K and S6A-B). While the spatial distributions of Sytox Green and DAPI were still highly similar between the two groups, *Cwf19l2*-SKO GV oocytes exhibited nearly twice AFI of Sytox Green than that of DAPI, and had a significantly higher Sytox Green-to-DAPI fluorescence intensity ratio (Figure 4L and S5C-E). This suggested that *Cwf19l2*-deficient GV oocytes retained too much abnormally unassigned RNA. Furthermore, we observed that *Cwf19l2*-SKO GV oocytes had lower OPP and HPG relative intensities compared to control oocytes (Figure 4M-N and S6F-G). As for splicing, SC35 was diffusely distributed throughout the nucleus in *Cwf19l2*-SKO GV oocytes, rather than aggregating into active speckles (Figure 4O). These observations indicated that maternal CWF19L2 ablation suppressed normal splicing and translation, while aberrantly elevating transcriptional activity in GV oocytes.

In summary, CWF19L2 depletion introduced latent defects at both the transcriptional and post-transcriptional levels during oogenesis, and these defects did not perturb morphological maturation or ovulatory capacity, but fully manifested upon ZGA, ultimately culminating in embryonic developmental arrest at the 2-cell stage.

### CWF19L2 loss elicits pronounced transcriptomic dysregulation

Given that CWF19L2 exhibited a dynamic splicing-dependent expression pattern within the nucleus, and interacted robustly with several core splicing factors, we hypothesized that CWF19L2 may functionally implicate in the regulation of AS in female germ cells and early embryos. To precisely and comprehensively test this conjecture, we isolated GV, MII oocytes, zygotes, late 2-cell embryos, and 48-hour post-fertilization embryos (normally progress to the 4-cell stage) from adult *Cwf19l2*-SKO and control mice, followed by Smart-seq2 analysis (Figure 5A). Transcriptomic analysis (*P* < 0.05 and |log_2_FoldChange| > 1.0, Supplementary Data 2) tracked the exact trajectory of molecular defects (Figure 5B). CWF19L2 depletion resulted in 722 identified differentially expressed genes (DEGs, 519 upregulated, 203 downregulated) in GV oocytes (Figure 5C), 1235 DEGs (871 upregulated, 364 downregulated) in MII oocytes (Figure S7A), and 1296 DEGs (930 upregulated, 366 downregulated) in zygotes (Figure S7B). Notably, the magnitude of transcriptional dysregulation peaked abruptly in late 2-cell embryos, 6963 DEGs (4475 upregulated, 2488 downregulated) (Figure 5E), aligning perfectly with the exact stage at which *Cwf19l2*-SKO embryos underwent developmental arrest. And there are 4421 DEGs (2732 upregulated, 1689 downregulated) in 48-hour post-fertilization embryos (Figure S7C).

**Figure 5.**
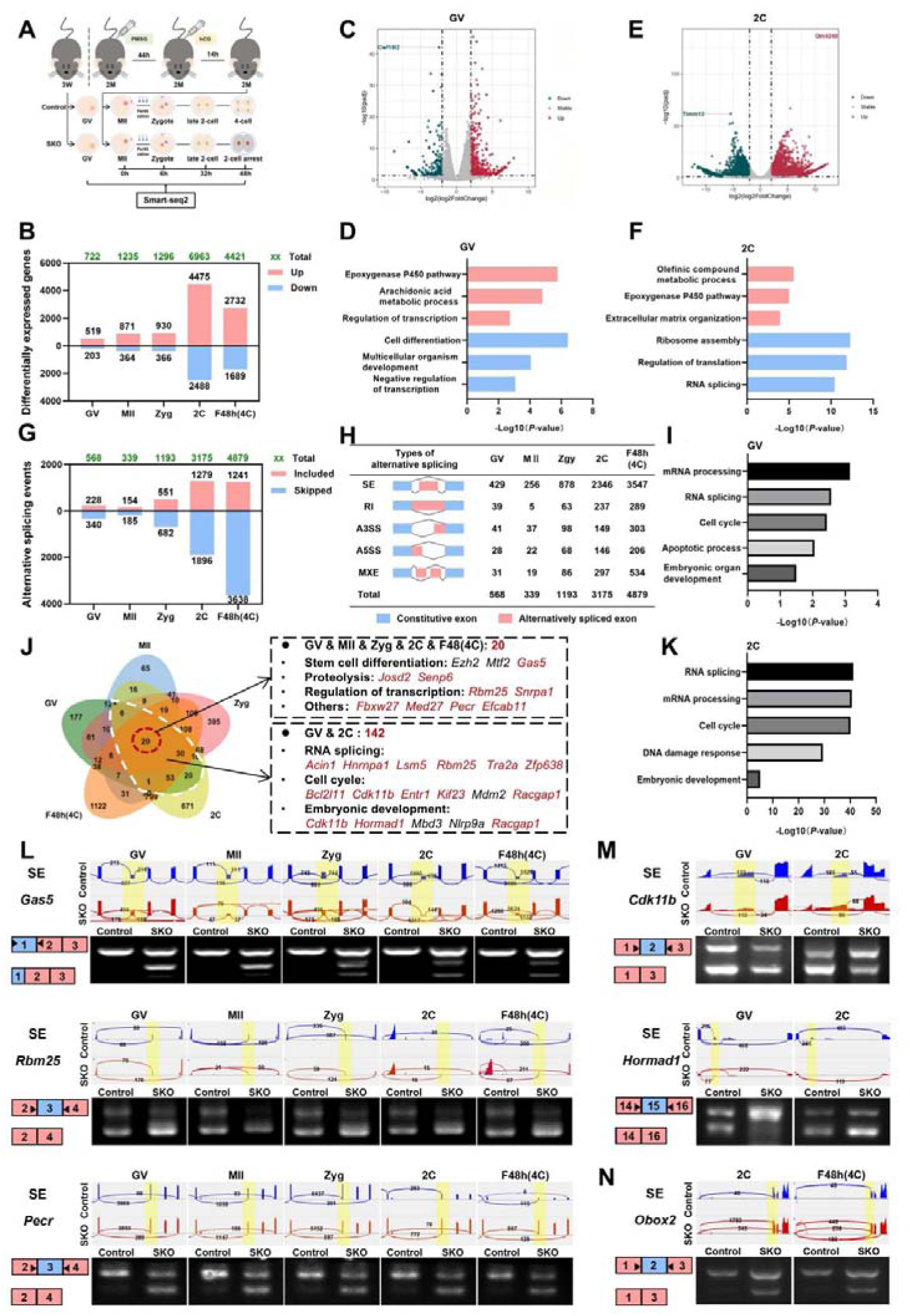
Transcriptomic landscape reveals widespread mRNA alternative splicing defects upon Cwf19l2 ablation **A.** Schematic diagram of sample collection for Smart-seq2 analysis. **B.** Number of DEGs in GV oocytes, MII oocytes, zygotes, 2-cell embryos, and 48 h post-fertilization embryos (F48h(4C)). Red indicates upregulated genes; blue indicates downregulated genes; and green indicates total number. **C.** Volcano plot of DEGs identified by Smart-seq2 in GV oocytes from Cwf19l2-SKO and control mice. Red represents upregulated genes, green represents downregulated genes, and gray represents genes with no significant difference. **D.** GO analysis of DEGs in GV oocytes. Red indicates upregulated genes; blue indicates downregulated genes. **E.** Volcano plot of DEGs in 2-cell embryos from Cwf19l2-SKO and control mice. **F.** GO analysis of DEGs in 2-cell embryos. **G.** Number of aberrant ASEs in GV oocytes, MII oocytes, zygotes, 2-cell embryos, and F48h(4C). Red indicates included events; blue indicates skipped events; and green indicates total number. **H.** Schematic diagram of five AS types and quantity of aberrant ASEs in GV oocytes, MII oocytes, zygotes, 2-cell embryos, and F48h(4C) embryos. The number of predicted ASEs in each category is indicated. **I.** GO analysis of ASEs in GV oocytes. **J.** Venn diagram showing the overlap of ASEs across 5-stage oocytes and early embryos from Cwf19l2-SKO and control mice. Red circles represent ASEs shared across all five stages; white circles represent ASEs common to GV oocytes and 2-cell embryos. GO analysis of common ASEs was listed on the right, and the successful validations were highlighted. **K.** GO analysis of ASEs in 2-cell embryos. **L.** Validation and validation of abnormal ASEs identified by Smart-seq2 in 5-stage oocytes and early embryos from Cwf19l2-SKO and control mice. Tracks from Integrative Genomics Viewer (IGV) for selected candidate genes are shown in the top panel, and differentially spliced parts are shaded. Schematics of ASEs are shown in the bottom-left panel. Gel images showing the RT-PCR analysis of ASEs of the changed splicing genes in the bottom-right panel. **M.** Validation and validation of abnormal ASEs identified by Smart-seq2 in GV oocytes and 2-cell embryos from Cwf19l2-SKO and control mice. **N.** Validation and validation of abnormal ASEs identified by Smart-seq2 in 2-cell and F48h(4C) embryos from Cwf19l2-SKO and control mice.

Subsequent Gene Ontology (GO) enrichment analysis decoded the functional consequences of these disruptions. In pre-fertilization oocytes (GV and MII), upregulated DEGs were enriched for transcriptional regulation, the peroxygenase P450 pathway, and lipid metabolic processes, whereas downregulated DEGs governed cell differentiation and developmental programs (Figure 5D and S7D). Post-fertilization, GO analysis of zygotes and embryos showed that upregulated genes were predominantly linked to apoptotic signaling, while downregulated genes were enriched in RNA processing, pre-mRNA splicing, and translational regulation (Figure 5F and S7E-F), providing a comprehensive molecular rationale for the severe EEA phenotypic defects observed previously (Figure 4D-O, S5-S6).

### Ablation of CWF19L2 provokes a catastrophic breakdown of alternative splicing homeostasis

Smart-seq2 data also uncovered the profound loss of AS fidelity in *Cwf19l2*-SKO oocytes and embryos, and we quantified aberrant alternative splicing events (ASEs) using stringent thresholds (*P* < 0.05 and |ΔPSI| > 10% (PSI: percent spliced in), Supplementary Data 3). In GV oocytes, 568 aberrant ASEs were identified, encompassing 429 skipped exons (SE), 39 retained introns (RI), 41 alternative 3’ splice sites (A3SS), 28 alternative 5’ splice sites (A5SS), and 31 mutually exclusive exons (MXE). This number declined to 339 in MII oocytes (256 SE, 5 RI, 37 A3SS, 22 A5SS, 19 MXE) before rebounding to 1193 in zygotes (878 SE, 63 RI, 98 A3SS, 68 A5SS, 86 MXE). Corresponding to the onset of ZGA, aberrant ASEs surged exponentially to 3,175 (2346 SE, 237 RI, 149 A3SS, 146 A5SS, 297 MXE) at the late 2-cell stage and further escalated to 4,879 (3547 SE, 289 RI, 303 A3SS, 206 A5SS, 534 MXE) in 48-hour post-fertilization embryos (Figure 5G-H). Notably, SE consistently emerged as the dominant aberrant ASE type across all stages. Chromosomal localization analysis of aberrant ASEs revealed highly conserved distribution patterns across all five stages (Figure S7G), without stage-specific enriched regions detected, indicating that CWF19L2 exerts a highly conserved regulatory role in AS.

Besides, we compared total aberrant ASEs with the number of alternative splicing genes (ASGs; genes harboring ≥1 ASE) to dissect the dynamics of splicing dysregulation. During oocyte transcriptional quiescence (GV to MII), the average number of aberrant ASEs per ASG decreased; however, following ZGA initiation, this metric exploded and even the most severe individual genes exhibited over 8 distinct aberrant (Figure S7H). GO analysis further showed that aberrant ASEs across all stages were consistently enriched in pathways critical for viability: RNA splicing, cell cycle control, and embryonic morphogenesis (Figure 5I, 5K, S7K-M). Together, our dynamic profiling highlights CWF19L2 as an indispensable safeguard against catastrophic splicing errors in oogenesis and embryogenesis.

### Persistent splicing errors induced by CWF19L2 ablation incapacitate core maternal and zygotic regulatory networks

Through systematic integration of ASEs across 5 developmental stages, we identified 20 core genes with consistent splicing disruption and further confirmed abnormal splicing for some of these genes (Figure 5J, 5L and S8A). For example, *Gas5*, the regulator of female germline stem cell proliferation and survival^[50, 51]^, produced aberrant transcripts lacking exon 1. *Rbm25*, a key mediator of embryonic stem cell pluripotency related to splicing control^[52, 53]^, exhibited exon 3 skipping. And exon 3 skipping was observed in *Pecr*, which was hypothesized to impair lipid metabolism^[54, 55]^, leading to energy homeostasis dysfunction in oocytes and embryos (Figure 5L). Furthermore, significant splicing abnormalities were observed in *Senp6* essential for embryonic survival^[56]^, *Snrpa1* governing gametogenesis^[57]^, *Efcab11* sustaining hyper prolificacy^[58]^, *Med27* directing early embryonic neurodevelopment^[59]^ and etc. (Figure S8A).

Given that maternal *Cwf19l2* depletion caused embryonic arrest at the 2-cell stage, and *Cwf19l2*-SKO GV oocytes already harbored splicing defects, we conducted a comparative analysis of aberrant ASEs between these two distinct stages, and identified 142 commonly mis-spliced genes. GO analysis of these common genes revealed enrichment in pathways related to RNA splicing (*Acin1*, *Hnrnpa1*, *Lsm5*, *Rbm25*, *Tra2a*), cell cycle progression *(Bcl2l11*, *Entr1*, *Kif23*), and embryonic morphogenesis (*Hormad1*) (Figure 5J). Over 80% of these ASEs were successfully validated across two stages (Figure 5M and S8B), emphasizing the embryos inherited maternal transcriptomic vulnerability. Additionally, we identified splicing abnormalities in *Obox* family genes, core regulators of ZGA^[60]^, in late 2-cell embryos and 48-hour post-fertilization embryos. The splicing defects included exon 2, exon 3, or combined exon 2/3 skipping in *Obox1*; exon 2 skipping in *Obox2;* exon 4 skipping in *Obox3*; and exon 2 skipping in *Obox8* (Figure 5N and S8C), which provided a direct mechanistic basis for the observed developmental blockade.

Together, these findings demonstrate that *Cwf19l2* depletion-induced AS dysregulation fundamentally disrupts cell cycle progression and embryonic development, and ultimately culminates in embryonic developmental arrest, highlighting that CWF19L2 enforces splicing fidelity through evolutionarily conserved regulatory mechanisms indispensable for early mammalian life.

### Splicing defects from CWF19L2 depletion cascades into widespread proteomic dysregulation

To validate whether these pervasive splicing errors compromise functional proteins, we performed ultra-low-input proteomic profiling on GV oocytes and 2-cell embryos derived from *Cwf19l2*-SKO and control mice (*P* < 0.05, Supplementary Data 4). The resulting proteomic landscape closely mirrored the transcriptomic breakdown. In GV oocytes, we identified 1087 differentially expressed proteins (DEPs), consisting of 400 upregulated and 687 downregulated candidates; in 2-cell embryos, 889 DEPs were detected, with 440 upregulated and 449 downregulated (Figure S9A). GO enrichment analysis further corroborated the consistency between proteomic and transcriptomic datasets. In GV oocytes, upregulated DEPs were primarily enriched in cholesterol metabolic process, apoptotic process and regulation of transcription, while downregulated DEPs were also associated with chromatin remodeling, cell differentiation, embryonic development and regulation of cell cycle (Figure S9B). This functional dichotomy persisted in 2-cell embryos, where upregulated proteins clustered in cellular response to lipid, and apoptotic process, and downregulated proteins were linked to gene expression, regulation of cell cycle, regulation of transcription, RNA splicing and regulation of translation (Figure S9C). Crucially, proteins encoded by genes with confirmed ASEs revealed a predominant downregulation trend across both stages (Figure S9D-E). Collectively, these multi-omic findings firmly establish that CWF19L2 ablation derails the precise splicing of key regulatory transcripts, which in turn triggers a widespread loss of essential proteins and ultimately precipitates the lethal developmental arrest observed in mutant mice.

### CWF19L2 directly orchestrates pre-mRNA splicing in oocytes and embryos

To dissect the molecular mechanism underlying CWF19L2-mediated AS regulation, we performed high-resolution linear amplification of complementary DNA ends and sequencing (LACE-seq) in GV oocytes and 2-cell embryos. LACE-seq analysis identified 859 binding peaks corresponding to 585 target genes in GV oocytes, and 3449 binding peaks across 1535 target genes were detected in 2-cell embryos (*P* < 0.05 and |log_2_FoldChange| > 1.0, Figure 6A–6B, Supplementary Data 5). Binding site annotation showed a preferential localization to intergenic and intronic regions of protein-coding genes in both stages (Figure 6A). Furthermore, motif enrichment analysis via the HOMER algorithm further delineated a conserved CWF19L2 consensus motif GGAASAGC (S=C or G) across oocytes and embryos (Figure 6C). The intrinsic RNA-binding capacity of CWF19L2 was validated in vivo by LACE-qPCR, which confirmed direct binding to key target transcripts, including *Fbxw27*, *Josd2*, *Tma16* and *Tpr* in both oocytes and embryos (Figure 6D-E).

**Figure 6.**
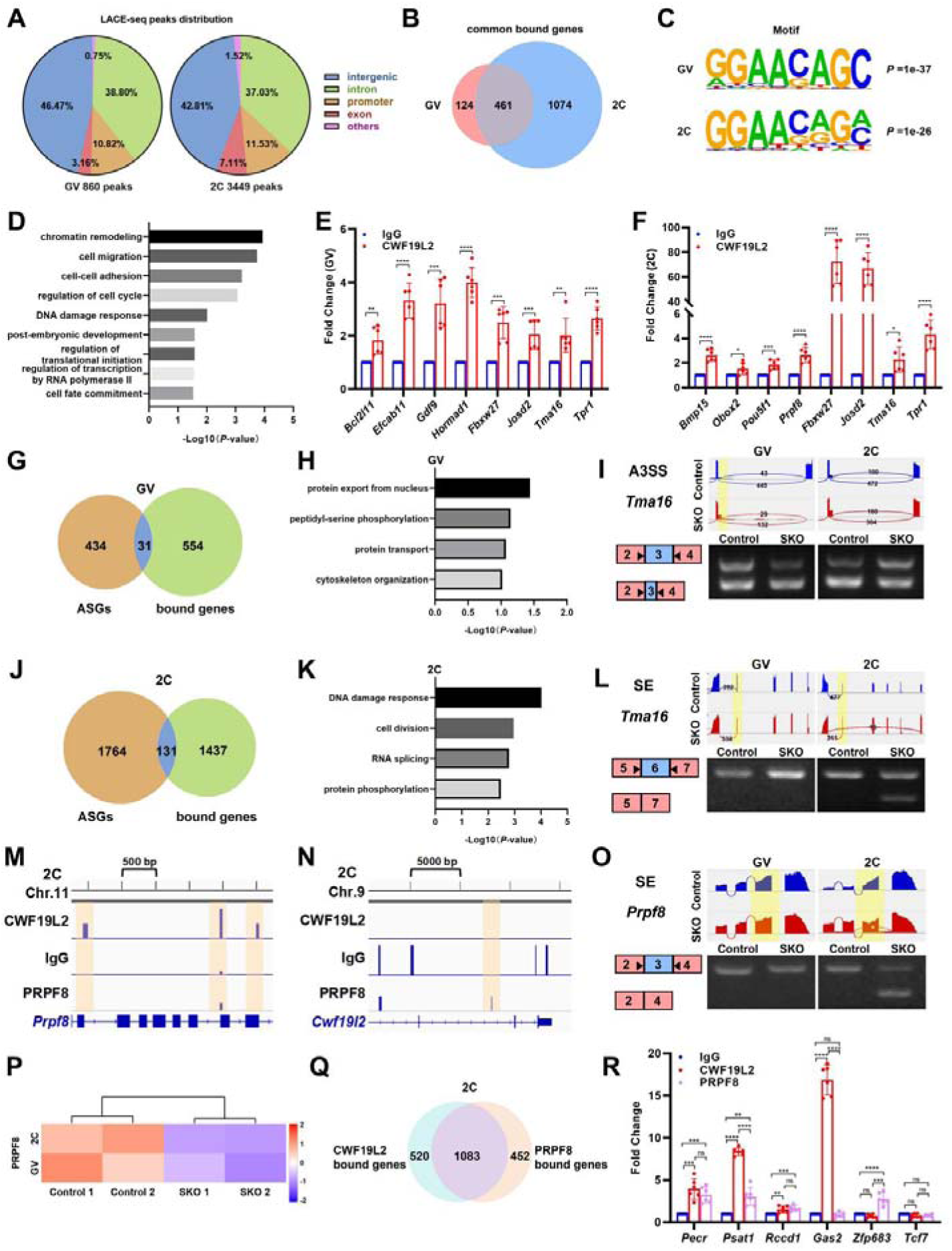
CWF19L2 directly regulates alternative splicing and indirectly controls splicing programs via modulation of splicing factors **A.** Pie chart showing the distribution of CWF19L2 binding locations in the genome from LACE-seq in GV oocytes and 2-cell embryos isolated from Cwf19l2-SKO and control mice. **B.** Venn diagram showing the common genes between CWF19L2-bound genes in GV oocytes and 2-cell embryos. **C.** HOMER de novo motif analysis of CWF19L2 binding peaks based on the LACE-seq that shows the identified motif. **D.** GO analysis of common bound genes between GV oocytes and 2-cell embryos. **E.** LACE-qPCR analysis of selected CWF19L2-bound genes in GV oocytes, using anti-CWF19L2 antibody and control IgG. Data are shown as mean ± SD, n = 3, **P < 0.01, ***P < 0.001, ****P < 0.0001. **F.** LACE-qPCR analysis of selected CWF19L2-bound genes in 2-cell embryos, using anti-CWF19L2 antibody and control IgG. Data are shown as mean ± SD, nn = 3, *P < 0.05, ***P < 0.001, ****P < 0.0001. **G.** Venn diagram showing the common genes between alternative splicing genes (ASGs) and CWF19L2-bound genes in GV oocytes. **H.** GO analysis of common genes between ASGs and CWF19L2-bound genes in GV oocytes. **I.** Visualization and validation of Tma16 abnormal A3SS in GV oocytes and 2-cell embryos from Cwf19l2-SKO and control mice. **J.** Venn diagram showing the common genes between alternative splicing genes (ASGs) and CWF19L2-bound genes in 2-cell embryos. **K.** GO analysis of common genes between ASGs and CWF19L2-bound genes in 2-cell embryos. **L.** Visualization and validation of Tma16 abnormal SE in GV oocytes and 2-cell embryos from Cwf19l2-SKO and control mice. **M-N.** Genome browser tracks showing LACE-seq binding peak distributions of CWF19L2, PRPF8, and IgG in Prpf8 locus (M) and Cwf19l2 locus (N) in 2-cell embryos. **O.** Visualization and validation of Prpf8 abnormal ASEs in GV oocytes and 2-cell embryos from Cwf19l2-SKO and control mice. **P.** Heatmap of protein expression for PRPF8 in GV oocytes and 2-cell embryos. **Q.** Venn diagram showing the common genes between CWF19L2 and PRPF8-bound genes in 2-cell embryos. **R.** LACE-qPCR analysis of selected mRNAs immunoprecipitated by anti-CWF19L2 and anti-PRPF8 antibodies and control IgG in 2 cell embryos. Data are presented as the mean ± SD, n = 3, ns: not significant, **P < 0.01, ***P < 0.001, ****P < 0.0001.

Cross-stage comparison revealed substantial overlap of CWF19L2 target genes between oocytes and embryos. Nearly 80% of target genes in GV oocytes (481/585) were also identified in 2-cell embryos (Figure 6B), indicating conserved mRNA targets across developmental stages. GO analysis of these shared genes linked them predominantly to chromatin remodeling, regulation of cell cycle, post-embryonic development and cell fate commitment, which were pathways crucial for cellular homeostasis and developmental progression (Figure 6F).

To link RNA-binding activity of CWF19L2 to its AS regulation, we integrated LACE-seq-derived target genes with ASEs identified by Smart-seq2. This multi-omic integration pinpointed 31 and 131 common genes in GV oocytes and 2-cell embryos, respectively (Figure 6G and 6J). GO analysis of these common direct targets in GV oocytes mainly associated with protein transport, whereas common genes in 2-cell embryos converged heavily on pathways essential for the DNA damage response and RNA splicing (Figure 6H and 6K). Strikingly, while transcripts such as *Tma16* and *Tpr* were bound by CWF19L2 at both stages, their *Cwf19l2* depletion-induced splicing defects manifested in a highly stage-specific manner. For *Tma16*, A3SS of exon 3 preferentially occurred in both GV oocytes and 2-cell embryos of control mice (Figure 6I), whereas exon 6 skipping emerged exclusively in the 2-cell embryo of *Cwf19l2*-SKO mice (Figure 6L). For *Tpr*, intron 42 retention was detected in both stages of *Cwf19l2*-SKO mice but exhibited more diverse retention in 2-cell embryos (Figure S10A), while exon 3 skipping was uniquely restricted to 2-cell embryos of *Cwf19l2*-SKO mice (Figure S10B). These stage-distinct splicing alterations demonstrate the dynamic, context-dependent nature of CWF19L2-mediated pre-mRNA splicing during MZT.

### CWF19L2 amplifies its splicing regulatory network via PRPF8

PRPF8 was also identified as a prominent target of CWF19L2 in our LACE-seq dataset, with close interaction have validated experimentally (Figure 6E). Track visualization revealed robust CWF19L2 occupancy at *Prpf8* locus in 2-cell embryos (Figure 6M). A similar binding pattern was observed in GV oocytes (Figure S10C), albeit with weaker signal intensity that precluded reliable detection by LACE-qPCR. Concomitantly, *Prpf8* exhibited exon 3 skipping in *Cwf19l2*-SKO 2-cell embryos (Figure 6O), and its protein expression was markedly reduced in both GV and 2-cell stages upon CWF19L2 depletion (Figure 6P). Therefore, we hypothesized that PRPF8 acts as a critical downstream mediator, indirectly amplifying CWF19L2-dependent splicing regulation.

To test this hypothesis, we used anti-PRPF8 antibody to perform LACE-seq in GV oocytes and 2-cell embryos. The track visualization showed negligible PRPF8 binding in *Cwf19l2* locus at either stage (Figure 6N and S10D), establishing a unidirectional regulatory axis. Comparative analysis of target genes in 2-cell embryos further revealed that approximately 70% of genes were co-regulated by both CWF19L2 and PRPF8 (e.g. *Pecr*, *Psat1*, *Rccd1*) (Figure 6Q-R). Crucially, the aberrant splicing of these shared targets was consistently detected in *Cwf19l2*-SKO 2-cell embryos (Figure 5L, S8B). In contrast, a subset of genes (like *Gas2*) was preferentially regulated by CWF19L2, while others (like *Zfp638)* were dominated by PRPF8 control (Figure 6R). This extensive regulatory overlap was also evident during oogenesis, where over half of the CWF19L2 targets in GV oocytes were shared with PRPF8 (Figure S10E). Taken together, our data establish that CWF19L2 modulates pre-mRNA alternative splicing through two distinct strategies, direct binding and indirect regulation mediated by PRPF8, to ensure cellular homeostasis during oocyte and early embryonic development.

### CWF19L2 deletion-induced early embryonic arrest can be partially ameliorated

To define the functional requirement of *Cwf19l2* during embryogenesis, we performed rescue experiments in maternal *Cwf19l2*-deficient embryos using mRNA microinjection. MII oocytes from *Cwf19l2*-SKO and control mice were fertilized with wild-type sperm, and viable zygotes were microinjected with mCherry-tagged *Cwf19l2* mRNA 6 hours post-fertilization (Figure 7A). Immunofluorescence staining confirmed robust expression of this protein, validating exogenous mRNA functionality (Figure 7B). Exogenous *Cwf19l2* mRNA supplementation enabled over 60% of maternal *Cwf19l2*-deficient zygotes to bypass the typical 2-cell arrest and proceed to later stages. However, despite resolving 2-cell arrest, most rescued embryos exhibited profound developmental delay; only one embryo reached the blastocyst stage by 96 hours post-fertilization (Figure 7C-D and S11A). Notably, microinjection of *Cwf19l2* mRNA into wild-type zygotes had no impact on embryonic development (Figure 7C and S11A-B).

**Figure 7.**
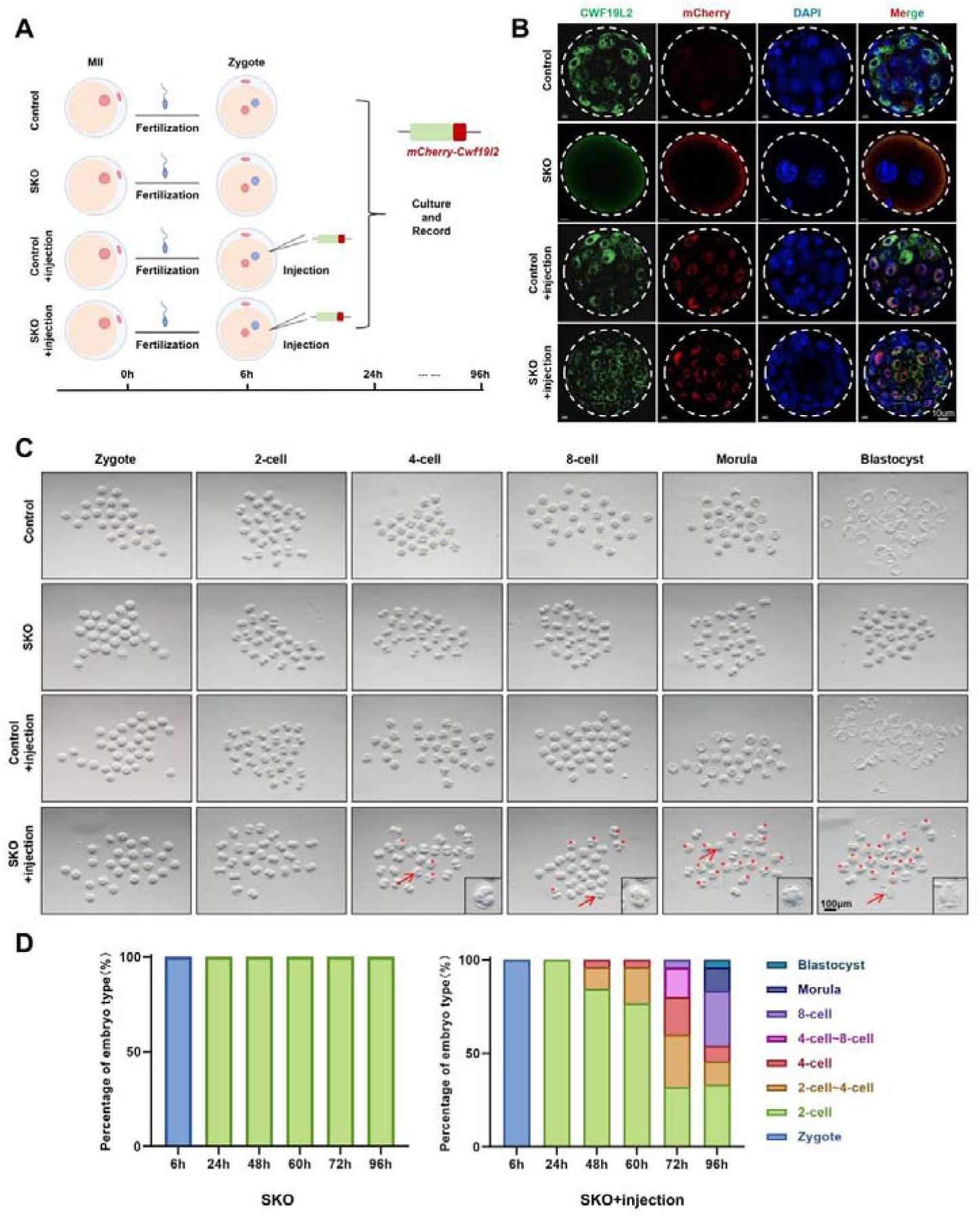
Exogenous Cwf19l2 partially rescues the 2-cell block phenotype of Cwf19l2-deficient embryos **A.** Schematic of the procedure for the rescue experiment. **B.** Co-immunofluorescence staining of CWF19L2 (green) with mCherry (red) in the most developed embryos from Cwf19l2-SKO and control mice with or without injection at 96h. DNA was stained with DAPI. Scale bars = 10 µm. **C.** Representative images of pre-implantation embryos at different stages derived from adult Cwf19l2-SKO and control mice with or without injection. Asterisks: embryos that have developed beyond the 2-cell stage. Arrows: the most developed embryos in the current time point are enlarged in the lower right corner. **D.** The percentage composition of different embryo types in adult Cwf19l2-SKO mice without or with injection at 6, 24, 48, 60, 72, and 96h after fertilization.

Taken together, these data demonstrate that post-fertilization delivery of exogenous *Cwf19l2* can partially mitigate the early embryonic arrest induced by maternal depletion. Crucially, the inability to fully restore normal developmental kinetics strongly implies that the latent transcriptomic and splicing damage accumulated during *Cwf19l2*-deficient oogenesis is largely irreversible by the zygotic stage, further cementing the indispensable role of CWF19L2 prior to fertilization.

## Discussion

AS is a fundamental post-transcriptional mechanism driving gene expression dynamics during gametogenesis and early embryogenesis^[23, 61]^. However, the specific splicing factors and molecular networks governing AS in female germ cells remain largely elusive^[24]^. Our study positions CWF19L2 as an indispensable orchestrator of AS through diverse mechanisms during MZT (Figure 8), thereby advancing our understanding of the splicing regulation in mammalian reproduction.

**Figure 8.**
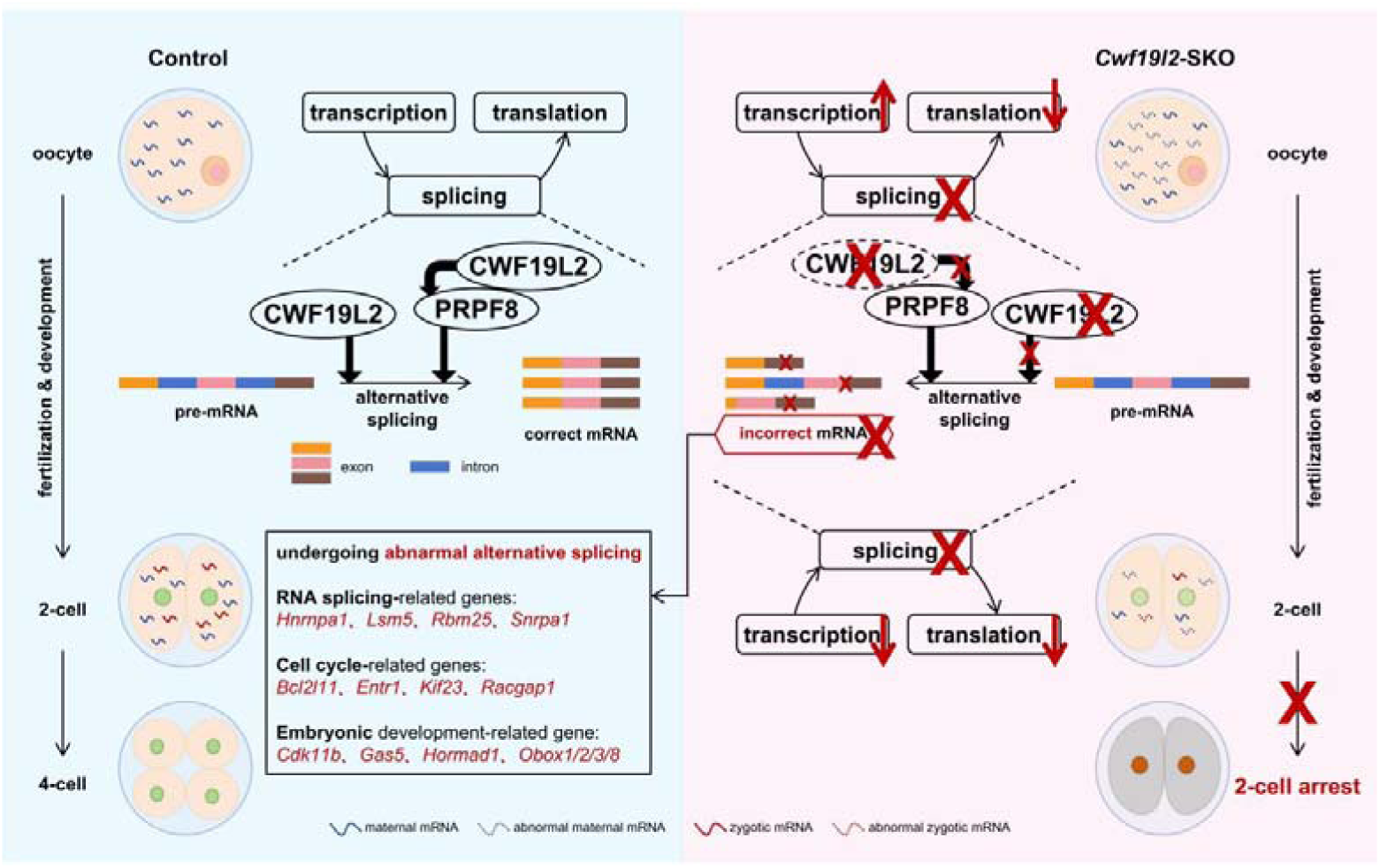
Schematic model of CWF19L2-mediated AS regulation in oogenesis and early embryogenesis. Schematic model showing that CWF19L2 directly binds to target genes to coordinate the proper AS of mRNA transcripts and that CWF19L2 controls key splicing factors like PRPF8 to indirectly amplify its role in splicing regulation during oogenesis and early embryogenesis.

### Evolutionary conservation and sexual dimorphism of CWF19L2 in gametogenesis

CWF19L2 exhibits profound evolutionary conservation by functionally integrating into the core spliceosome^[39, 40]^. Consistent with prior observations in spermatocytes, we validated its robust interactions with PRPF8, PRPF43, and SLU7 in HEK-293T cells. Furthermore, our domain-mapping analyses explicitly demonstrated that the CWFJ domain of CWF19L2 physically interacts with three core domains of PRPF8, substantially advancing our structural and mechanistic understanding of this regulatory module. While material scarcity and strict animal welfare constraints precluded conventional Co-IP in vivo, we deployed immunofluorescence and PLA to bridge this technical gap. We confirmed CWF19L2 robustly colocalizes with core spliceosomal proteins in oocytes and embryos. Most critically, PLA resolved the ultra-structural spatial proximity (< 40 nm) between CWF19L2 and PRPF8, providing compelling in situ evidence to support their direct physical interaction in females. Although future breakthroughs in ultra-low-input biochemical profiling will be required to physically confirm this interaction, our current multi-dimensional evidence establishes a solid mechanistic foundation. Together, this conserved structural integration provides the basal machinery required for stable splicing cycles, ensuring that CWF19L2 orchestrates spliceosome assembly and precise pre-mRNA splicing in oocytes and embryos, as crucially as it does in spermatogenesis.

A striking paradigm emerging from our study is the profound sexual dimorphism elicited by *Cwf19l2* ablation. While CWF19L2 is absolutely indispensable for fertility in both sexes, the spatiotemporal manifestations of its deletion are drastically uncoupled. In males, *Cwf19l2* deficiency triggers an immediate, catastrophic collapse of spermatogenesis. Conversely, mutant females execute morphologically normal oogenesis but suffer absolute embryonic developmental arrest at 2-cell stage. Spermatogenesis is characterized by rapid, continuous mitotic and meiotic proliferation, rendering male germ cells acutely and immediately vulnerable to splicing defects^[62, 63]^. In stark contrast, dictyate oocytes endure prolonged transcriptional quiescence and prioritize the accumulation of maternal reserves^[64, 65]^. This unique developmental pacing allows oocytes to morphologically tolerate latent transcriptomic scarring, which only becomes lethal when strictly zygotic de novo RNA processing is demanded during the MZT. Ultimately, this sex-specific divergence highlights how a highly conserved basal splicing machinery is uniquely wired by divergent developmental programs to meet the distinct metabolic and kinetic demands of male versus female reproduction.

This macroscopic phenotypic divergence is fundamentally mirrored at the molecular level. Although similar GO analysis of DEGs and ASEs were observed in the transcriptome of male and female gametes between *Cwf19l2*-SKO and control mice, the results exhibited minimal overlap (45 common DEGs and 58 common ASGs between male spermatocytes and GV-stage oocytes, and 72 common DEGs and 43 common ASGs between spermatocytes and MD-stage oocytes). As for, we hypothesized that CWF19L2 does not impose a rigid, universal splicing program. Instead, it functions as a highly adaptable orchestrator that services the intrinsically distinct RNA process environments of each sex. Besides, the differences may also be from the intrinsic gene expression profiles between male and female mice.

### The dual-phase vicious cycle of transcription-splicing-translation uncoupling induced by CWF19L2 deletion

Successful MZT required exquisite coordination of transcription, splicing, and translation to orchestrate proper genes expression. Our findings propose a stage-specific, dual-phase pathogenic vicious cycle wherein *Cwf19l2* depletion fundamentally uncouples this regulatory axis across oogenesis and embryogenesis.

During the GV stage, this uncoupling manifests as a compensatory vicious cycle. We postulate that the initial paralysis of the splicing machinery starves the translational apparatus of mature mRNA templates, accompanied by insufficient maternal substances. In a futile compensatory response, the oocyte hyper-activates transcription to generate surplus pre-mRNA templates, which merely exacerbates the toxic accumulation of unprocessed RNA. However, early embryos suffer from a source-depleted vicious cycle. Upon ZGA, a residual burst of early embryonic transcription yields only limited nascent RNA substrates. Because the splicing machinery has completely collapsed by this stage, these scarce transcripts cannot be properly processed. This absolute void of mature mRNA templates ultimately precipitates a catastrophic translational crash, sealing the fate of embryonic lethality. Thus, the 2-cell developmental block is not strictly an intrinsic embryonic failure, but rather the ultimate, irreversible manifestation of profoundly compromised oocyte quality, fitting perfectly within the clinical etiology of OECD.

Remarkably, despite established defects in RNA processing and protein synthesis*, Cwf19l2*-deficient oocytes could still complete meiosis and achieve successful fertilization. This delayed penetrance likely arises because oocytes possess an intrinsically high fault tolerance to ensure reproduction; as long as fundamental meiotic checkpoints are satisfied, they exert minimal quality control over the specific maternal determinants required for later embryogenesis. Temporally, since the *Stra8-Cre* exerted between E12.5 and E16.5, when maternal substances supporting embryonic development may have already been preferentially synthesized and stockpiled.

### CWF19L2 divers a bipartite and highly dynamic regulatory paradigm

Mechanistically, we postulate that CWF19L2 exerts its regulatory influence through a sophisticated bipartite mechanism. It not only directly binds and executes the splicing of critical transcripts but also amplifies its regulatory function indirectly through PRPF8. While these dual approaches parallel the regulatory paradigm observed in spermatogonia, the downstream molecular targets diverge significantly, indicating that the full expanse of the CWF19L2-driven splicing network warrants further elucidation.

Moreover, the stage-specific splicing alterations observed in conserved targets confirm the highly dynamic nature of CWF19L2. It continuously recalibrates its splicing output to accommodate the rapidly evolving synthesis requirements from oocytes and embryos, rather than being static. This dynamic tuning ensures RNA homeostasis and accounts for the dynamic shifts in shared target genes between GV and 2-cell stages. Unraveling the precise molecular determinants that enable CWF19L2 to orchestrate such stage-specific isoform expression will be a critical direction for future studies.

### Irreversibility of maternal molecular scarring

The inability of post-fertilization exogenous *Cwf19l2* to fully restore normal development provides critical insight into the ontogeny of MZT failure. While zygotic intervention can bypass the immediate 2-cell block, it cannot overwrite the deep-seated maternal factor deficit and widespread transcriptional and post-transcriptional defects accumulated in oocytes. Whether injecting relevant downstream targets or performing injections in the earlier oocytes could achieve more successful rescue remains to be further explored. This temporal partition emphasizes that the therapeutic window for addressing certain etiologies of female infertility could be shifted upstream to the oocyte maturation phase.

In conclusion, our research establishes CWF19L2 as a maternal splicing factor that safeguards the integrity of oogenesis and embryogenesis, and has a dual influence on alternative splicing, encompassing both direct modulation of essential genes like *Obox2* and indirect regulation mediated by PRPF8. This study fundamentally advances our understanding of the AS during gametogenesis and early embryonic development, and provides new perspectives for genetic and molecular basis of infertility and also offers novel clinical therapeutic ideas for the treatment of OECD in human reproductive medicine.

## Methods

### Mice and genotyping

C57BL/6J-background *Cwf19l2^flox/+^* mice (Cyagen Biosciences) were generated by ESC-mediated targeted modification and blastocyst microinjection, with two loxP sites flanking the 6th exon of *Cwf19l2* in ESCs. C57BL/6J-background *Stra8-GFPCre* mice were a generous gift from Prof. Ming-han Tong (Center for Excellence in Molecular Cell Science, Chinese Academy of Sciences) ^[48, 66]^.

For strain generation, *Cwf19l2^flox/flox^* mice were mated with *Stra8-GFPCre* mice to generate the C*wf19l2^flox/+^*; *Stra8-GFPCre* mice, and then these mice were bred with *Cwf19l2^flox/+^ or Cwf19l2^flox/flox^*mice to obtain *Cwf19l2^flox/flox^*; *Stra8-GFPCre* (*Cwf19l2*-SKO or SKO) mice.

All mice were maintained under standard conditions (21-22 °C, 12 h: 12 h light/dark cycle, ad libitum water and food access). Genotyping was performed via PCR on genomic DNA extracted from PD5 mouse tail tissues, with primer sequences in Supplementary Data 6. All experiments were approved by the Regional Ethics Committee of Shandong University.

### Fertility test

For female fertility testes, individual 6-week-old *Cwf19l2*-SKO females were cohoused with a single 12-week-old WT C57BL/6J male in a cage, and these tested was conducted continuously for at least 6 months. Females were inspected daily for vaginal plug formation and tracked pregnancy outcomes.

### Histological analysis

Following euthanasia, ovaries were fixed overnight at 4Din Bouin’s solution (Scientific Phygene, PH0976). Tissues were dehydrated, embedded in paraffin, and sectioned at 5 μm. After deparaffinization, sections were stained with hematoxylin following a standard protocol. Images were acquired using an Olympus BX53 fluorescence microscope.

### Oocyte collection

For NSN-GV oocytes, ovaries isolated from 2-week-old female mice were digested in pre-warmed digestion medium for 15 min at 37Dunder 5% CO_2_ and 5% O_2_, and then NSN-GV oocytes were collected and washed thoroughly to remove residual granulosa cells. SN-GV oocytes were collected from 3-week-old female mice, which were intraperitoneally injected with 5 IU PMSG. Forty-six hours post-injection, oocytes were harvested as above. MII oocytes were collected from 6-week-old mice, which were initially primed with PMSG (Ningbo, China) for 46 h followed by priming with hCG (Ningbo, China) for 14 h. Cumulus-oocyte complexes (COCs) from oviducts were released into pre-warmed hyaluronidase solution and incubated at 37Dfor 5 minutes to remove cumulus cells. Denuded oocytes were washed and maintained under appropriate culture conditions for subsequent.

### In vitro fertilization and embryo culture

The ovulated MII oocytes were inseminated with normal sperm isolated from adult male mice in pre-equilibrated IVF medium (Vitrolife). 6 hours post-insemination, zygotes with pronuclear formation were transferred to G1 medium (Vitrolife) under appropriate culture conditions for further experiments. Embryonic development was monitored daily: the attainment of the early 2-late 2-, 4-, 8-cell, morula, and blastocyst stages were recorded on 24, 36, 48, 60, 72, and 96 hours post-insemination.

### Microinjection

Microinjection was performed using a Nikon ECLIPSE Ti2 microinjector. *Cwf19l2* plasmids were synthesized by Miaoling Biotechnology, and mRNA was generated via in vitro transcription. A 10 pL volume of mRNA solution (500 µg/mL) was injected into each zygote. Post-injection, zygotes were transferred to G1-plus medium and cultured overnight in an incubator at 37 D with 5% COD and 5% OD to support subsequent development.

### Immunofluorescence

Oocytes and embryos were fixed in 4% paraformaldehyde (Solarbio, P1110) for 30 min at room temperature (RT), washed three times in wash buffer (0.1% SDS in PBS), permeabilized in 0.1% Triton X-100 in PBS for 15 min at RT, washed again as above, and blocked in 5% BSA for 30 min at RT. Then, the samples were incubated with primary antibody diluted in 5% BSA overnight at 4 °C. After washing three times in wash buffer, the samples were incubated with secondary antibody diluted in PBS for 1 h at RT in the dark, washed again as above, and mounted with DAPI Fluoroshield mounting medium (Abcam, ab104139). Images were acquired using an Andor Dragonfly spinning-disc confocal microscope controlled by Fusion software, and were processed and evaluated by ImageJ. Antibody details are provided in Supplementary Data 7.

### Immunoblotting

Approximately 100 oocytes or 60 early embryos were washed three times in PBS, transferred to a tube with ≤3□µL residual volume, and lysed in 20□µL 1× SDS loading buffer (Beyotime, P0015L) by vortexing for 30Dseconds. Lysates were centrifuged at 15,000□rpm for 1□min, then denatured at 95□□ for 5□min.

HEK293T cells were collected and lysed in NP-40 Lysis Buffer (Beyotime, P0013F) with protease inhibitor (Roche, 04693132001). Following homogenization, extracts were incubated on ice for 20 min, and centrifuged at 4□at 15,000□rpm for 20 min. The supernatant was denatured with 5× SDS loading buffer, and denatured at 95□□for 5□min.

Equal total proteins were resolved on 10% SDS–PAGE gel (Invitrogen, NP0315) and transferred to PVDF membranes (Millipore, ISEQ00010). Membranes were blocked with 5% nonfat milk for 1 h at RT, incubated with primary antibodies overnight at 4°C, washed three times with TBST, and incubated with the secondary antibodies for 1 h at RT. Signals were detected by ClarityTM Western ECL Substrate (BioRad, 1705060). Antibody details are provided in Supplementary DataD7.

### Co-Immunoprecipitation

HEK 293T cells were cultured in 6-well plates and were co-transfected with 500 ng of Flag-tagged full-length or truncated *Cwf19l2* and Myc-tagged full-length or truncated *Prpf8* plasmids using HP DNA Transfection reagent (Roche, 06366546001). 48h later, cells were collected and processed into supernatant by the method in immunoblotting. The supernatant was incubated with primary antibodies and spun at 4 □overnight. Next day, the pre-cleaned magnetic protein A/G beads were added, and the mixture was spun at 4 □for 2h. The beads were washed with NP-40 Lysis Buffer with protease inhibitor three times and boiled in 2×SDS loading buffer for immunoblotting analyses.

### Proximity Ligation Assay (PLA)

Duolink^®^ PLA Multicolor Reagent Pack (Sigma-Aldrich, DUO96000) and Duolink^®^ PLA Multicolor Probemaker Kit - Red (Sigma-Aldrich, DUO96010) were utilized following the manufacturer’s instructions on GV oocytes, 2-cell embryos, and HEK 293T cells.

### qPCR and RT-PCR

cDNA was extracted from oocytes and embryos using FlysisAmp Cells-to-cDNA Kit (Vazyme, CL111-01) following the manufacturer’s instructions. qPCR reactions were performed in triplicate using SYBR Green Premix Pro Taq HS qPCR Kit (AG, 11701) on a LightCycler 96 system (Roche). Gapdh served as an internal control. Relative gene expression was calculated using the 2^-ΔΔCt^ method.

For RT-PCR, amplification is performed using 2×Es Taq MasterMix (Dye) (CWBIO, CW0690M) and specific primers under specific conditions. PCR products were separated on agarose gels, imaged, and quantified. All primer sequences are listed in Supplementary Data 6.

### SMART-seq2 and analysis

Single-cell RNA-seq libraries were prepared using the Discover-sc WTA Kit V2 (Vazyme, N711) for cell lysis, mRNA reverse transcription, and cDNA amplification. Amplified cDNA was fragmented and converted into sequencing libraries using the TruePrep Flexible DNA Library Prep Kit for Illumina (Vazyme, TD504). All steps were performed according to the manufacturer’s instructions. Libraries were sequenced by Novogene on an Illumina platform with 150 bp paired-end reads.

Raw paired-end reads were quality-checked with FastQC and trimmed using Trim Galore. The resulting clean reads were aligned to the mouse reference genome (mm10) using HISAT2 2 (version 2.0.5)^[67]^, and gene-level counts were quantified via featureCounts (version 1.5.0-p3)^[68]^. Expression levels were normalized to FPKM, and log-transformed data were used for sample correlation and principal component analysis. DEGs were performed with DESeq2. ASEs were identified via rMATS and visualized using rmats2sashimiplot. GO analysis was conducted using the clusterProfiler R package.

### LACE-seq and analysis

Linear amplification of cDNA ends and sequencing of oocytes and embryos was executed based on established protocols to identify specific RNA-protein interactions^[69]^. Initially, samples were subjected to UV-C irradiation to securely crosslink RNA to interacting proteins. Target complexes were then isolated via immunoprecipitation using specific antibodies conjugated to Protein A/G magnetic beads. Following cell lysis and DNA digestion, the immunoprecipitated RNA was fragmented using micrococcal nuclease. The RNA fragments subsequently underwent dephosphorylation before a 3’ cDNA linker was ligated utilizing T4 RNA Ligase 2. This ligation enabled first-strand reverse transcription, which was driven by a T7 primer and SuperScript II to synthesize the initial cDNA strand. To ensure library purity, residual primers were degraded using Exo I and RNase H. The newly synthesized cDNA was then captured and enriched using streptavidin C1 magnetic beads. Next, a second 3’ linker was ligated to this enriched cDNA pool via truncated T4 RNA Ligase 1, which paved the way for the primary round of PCR amplification. To achieve the linear amplification characteristic of LACE-seq, in vitro transcription was conducted over 24 hours using T7 polymerase, effectively multiplying the RNA targets. After digesting any leftover DNA, this amplified RNA was purified and utilized as a template for second-strand reverse transcription with a dedicated CLIP primer. A final round of PCR amplification solidified the sequencing library. To conclude the physical protocol, targeted amplicons strictly within the 130 to 400 base pair range were size-selected through agarose gel electrophoresis and purified. Single-end sequencing was performed on the Illumina HiSeq 2500 platform by Berry Genomics.

Raw sequencing reads were quality-filtered and trimmed for adapters, followed by alignment to the mouse reference genome using Bowtie software (v.1.2.3)^[70]^. Enriched binding clusters were identified using Piranha (http://smithlabresearch.org/software/piranha/, v.1.2.1) and annotated to local genomic features. Motif enrichment within peaks was evaluated via HOMER. Sample reproducibility was assessed with deepTools, while differential binding analysis was conducted using DiffBind and DESeq2. Differentially bound peaks were subsequently annotated to target genes for GO enrichment analysis.

### Statistical analysis

Quantitative experiments were based on at least three independent biological samples, and the data are presented as means ± standard deviation (SD). Statistical analyses were performed using GraphPad Prism 10.1.2 software (GraphPad, San Diego, CA, USA), employing a Student’s two-tailed *t*-test for pairwise comparisons. The data were considered significant when the *p*-value was less than 0.05.

## Supporting information

Sup2-Differentially expressed genes identified by Smart-seq2

Sup3-Abnormal alternative splicing events identified by Smart-seq2

Sup4-Proteomics results of GV oocytes and 2-cell embryos

Sup5-Target genes of CWF19L2 detected in LACE-seq

Sup6-Primer sequences are used in this study

Sup7-Antibodies are used in this study

## Supplementary information

Supplementary Data 1: **Figure S1-S11**.

Supplementary Data 2: **Table S1**. Differentially expressed genes identified by Smart-seq2.

Supplementary Data 3: **Table S2**. Abnormal alternative splicing events identified by Smart-seq2.

Supplementary Data 4: **Table S3**. Proteomics results of GV oocytes and 2-cell embryos.

Supplementary Data 5: **Table S4**. Target genes of CWF19L2 detected in LACE-seq.

Supplementary Data 6: **Table S5**. Primer sequences are used in this study.

Supplementary Data 7: **Table S6**. Antibodies are used in this study.

## Acknowledgments

We convey our sincere appreciation to all colleagues in the Chen Laboratory of Institute of Women, Children and Reproductive Health in SDU and Center for Reproductive Medicine of The First Affiliated Hospital of ZZU for their indispensable support. Further acknowledgment is owed to the Translational Medicine Core Facility of SDU for providing expert consultation and granting access to critical instrumentation. We also thank Prof. Heng-Yu Fan (Life Sciences Institute, Zhejiang University) for his insightful comments and guidance, which significantly improved this manuscript.

## Author Contributions

S.W. was involved in study design, conceptualization, the majority of experimentation, data collection, analysis, writing original draft, and review and editing. T.L. performed Smart-seq2 and LACE-seq data analysis. Y.C. provides technical guidance on LACE-seq. K.S., Z.W., and J.X. participated in animal husbandry, genotyping, and functional experiments. Q.Z., C.H., Y.S. and H.Z. participated in manuscript revision and facilitated the study design. T.H., Z.-J.C., H.L., Y.G., and H.Z. were responsible for study design, conceptualization, analysis, review, editing, supervision, project administration, resources, and funding acquisition. All authors have read and approved the final version of the manuscript.

