## Supplementary material for "CWF19L2 couples pre-mRNA alternative splicing with the maternal-to-zygotic transition to safeguard female fertility": Sup1-FigureS1-S11

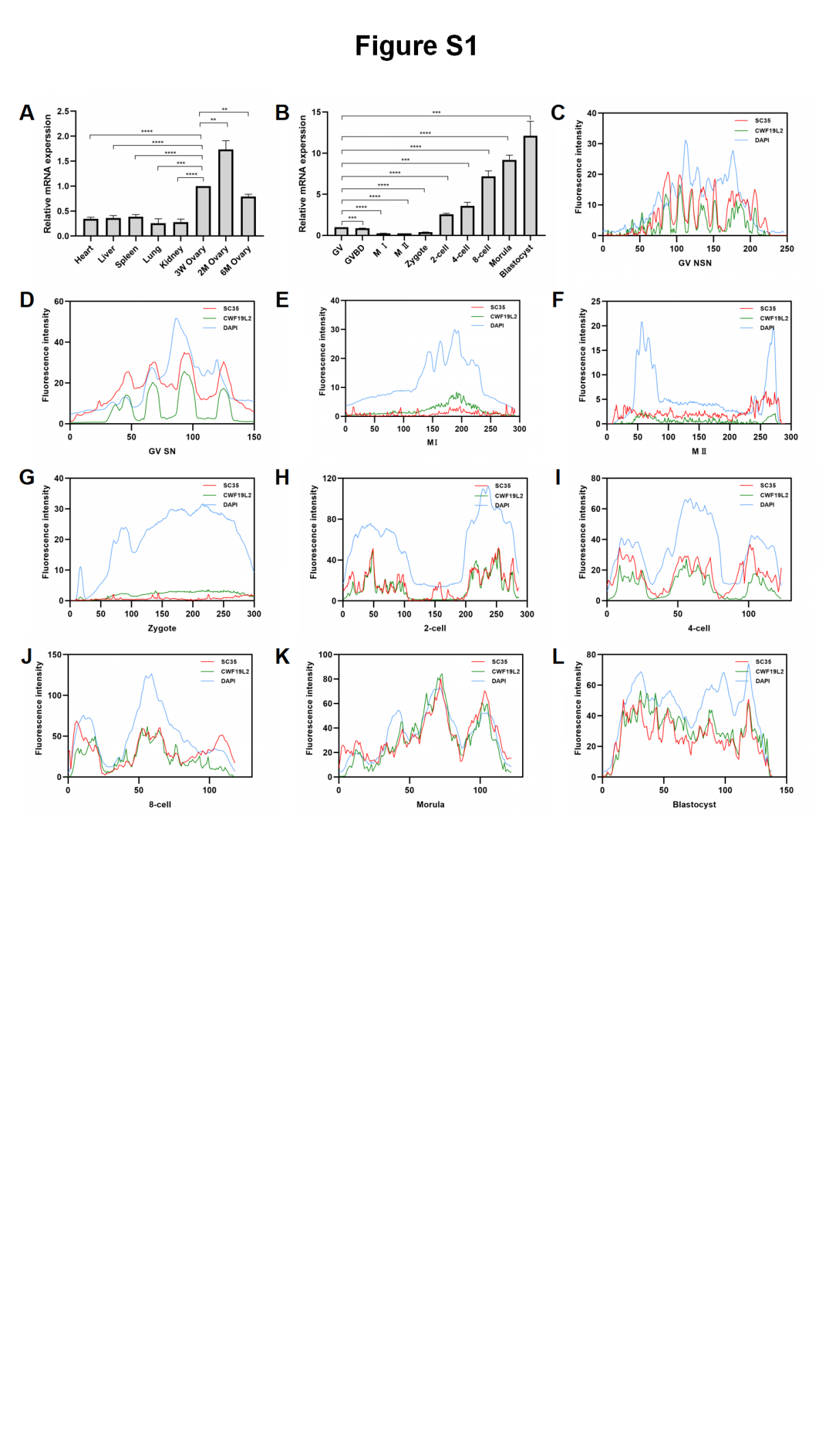

**Figure S1. Expression pattern of CWF19L2 during oogenesis and embryogenesis**

(A) QPCR analysis of the mRNA levels of *Cwf19l2* in multiple organs of adult WT mice. mRNA expression in ovary of 3-week-old (3W) mice was set as one. Data are presented as mean ± SD, n = 3,**:*P* < 0.01, ***:*P* < 0.001, ****:*P* < 0.0001.
(B) QPCR analysis of the mRNA levels of *Cwf19l2* at different stages of oocytes and embryos of adult WT mice. mRNA expression in germinal vesicle (GV) oocytes was set as one. Data are presented as mean ± SD, n = 3,***:*P* < 0.001, ****:*P* < 0.0001.

(C-L) Intensity profiles of SC35 (red), CWF19L2 (green), and DAPI (blue) in NSN GV, SN GV, MⅠ, MⅡ oocyte, zygote, 2-cell, 4-cell, 8-cell embryo, morula, and blastocyst respectively( Figure 1D) of *Cwf19l2*-SKO and control mice.

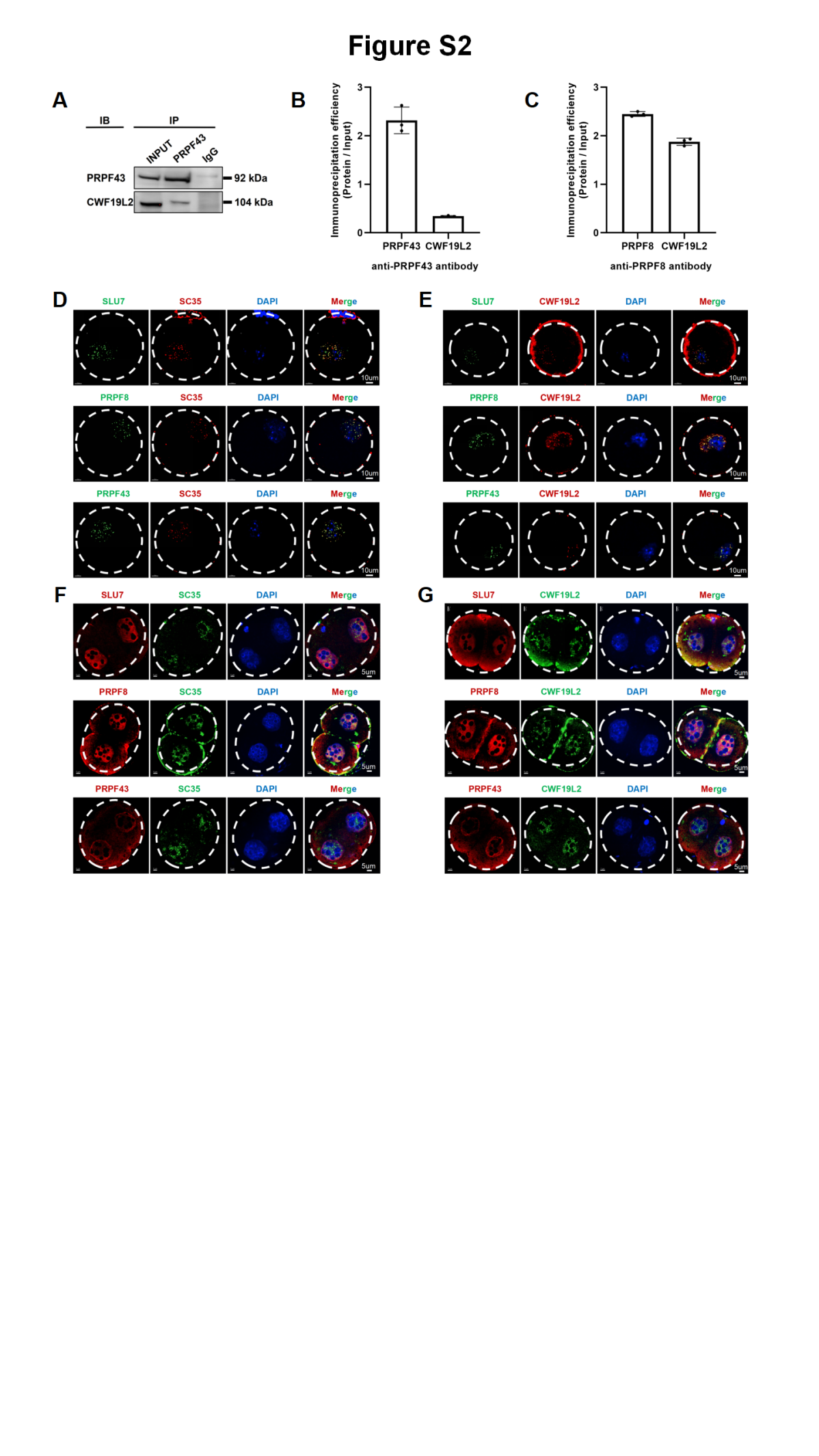

**Figure S2. Colocalization of CWF19L2 and splicing-related proteins in oocytes and embryos**

(A) Co-IP assays of the interaction between PRPF43 and CWF19L2 in HEK293T cells. IgG was used as the negative control.
(B) Relative immunoprecipitation efficiency of anti-PRPF43 antibody immunoprecipitated PRPF43 and CWF19L2 in sHEK293T cells.
(C) Relative immunoprecipitation efficiency of anti-PRPF8 antibody immunoprecipitated PRPF8 and CWF19L2 in sHEK293T cells.
(D) Co-immunofluorescence staining of nuclear speckles marker SC35 (red) with the SLU7, PRPF8 and PRPF43 (green) in GV oocytes of adult WT mice. DNA was stained with DAPI. Scale bars = 10 µm.
(E) Co-immunofluorescence staining of CWF19L2 (red) with the SLU7, PRPF8 and PRPF43 (green) in GV oocytes of adult WT mice. DNA was stained with DAPI. Scale bars = 10 µm.
(F) Co-immunofluorescence staining of nuclear speckles marker SC35 (green) with the SLU7, PRPF8 and PRPF43 (red) in 2-cell embryos of adult WT mice. DNA was stained with DAPI. Scale bars = 5 µm.
(G) Co-immunofluorescence staining of CWF19L2 (green) with the SLU7, PRPF8 and PRPF43 (red) in 2-cell embryos of adult WT mice. DNA was stained with DAPI. Scale bars = 5 µm.

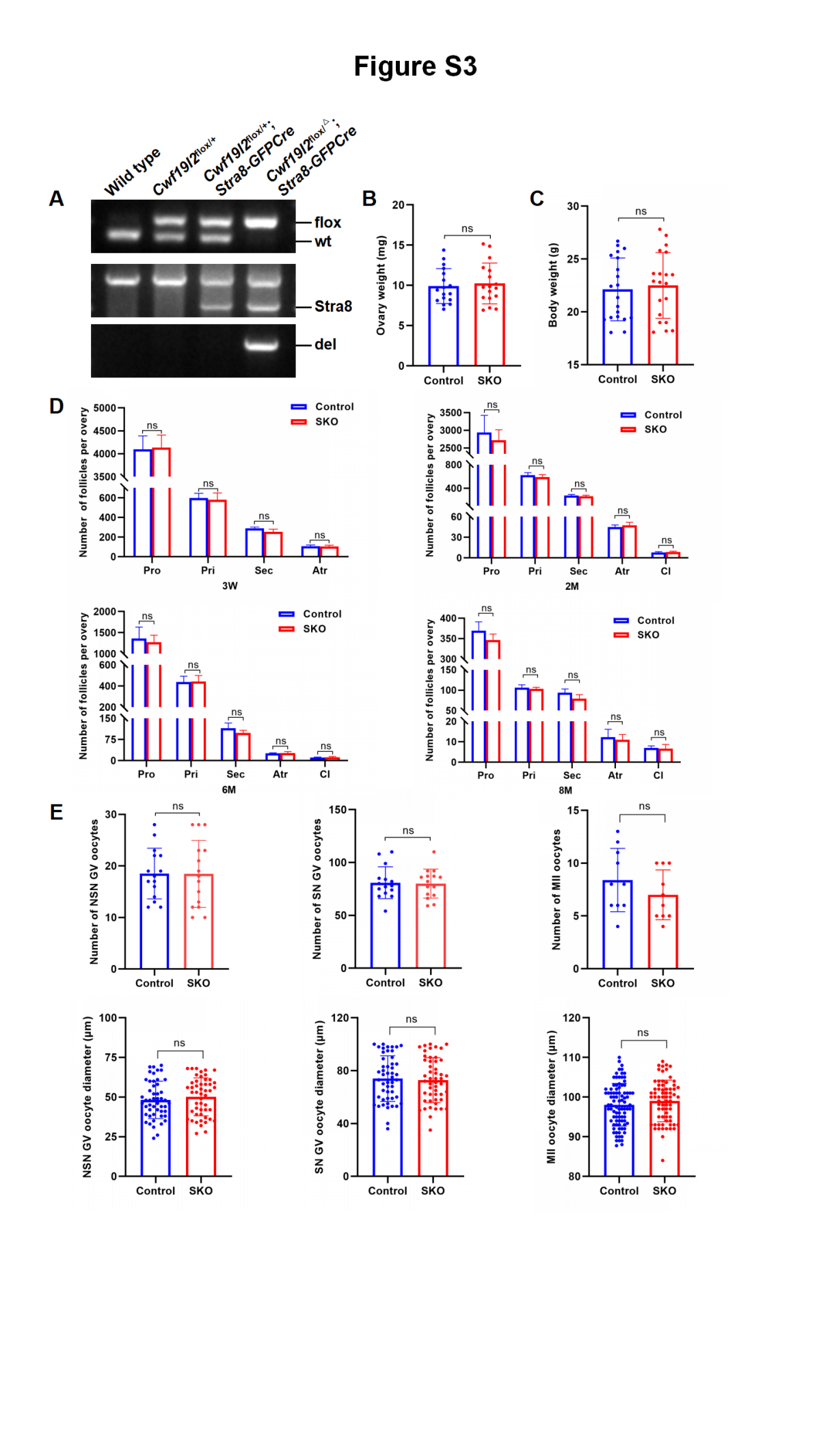

**Figure S3. CWF19L2 is not required for the morphological development of oogenesis in mice**

(A) PCR-based genotype of the PD5 mouse tails is shown.
(B) Ovary weight of adult *Cwf19l2*-SKO and control mice. Data are presented as the mean ± SD, n = 20, ns: not significant.
(C) Body weight of adult *Cwf19l2*-SKO and control mice. Data are presented as the mean ± SD, n = 20, ns: not significant.
(D) Quantification of follicles in overies of 3-week, 2-month, 6-month and 8-month *Cwf19l2*-SKO and control mice. Pro: primordial follicles; pri: primary follicles; Sec: secondary follicles; atr: antral follicles; CL: corpus luteum. Data are presented as mean ± SD, n=3, ns: not significant.
(E) Quantification of numbers (upper panel) and diameter (lower panel) of NSN GV, SN GV and MII oocytes obtained from *Cwf19l2*-SKO and control mice . Data are presented as mean ± SD, , ns: not significant.

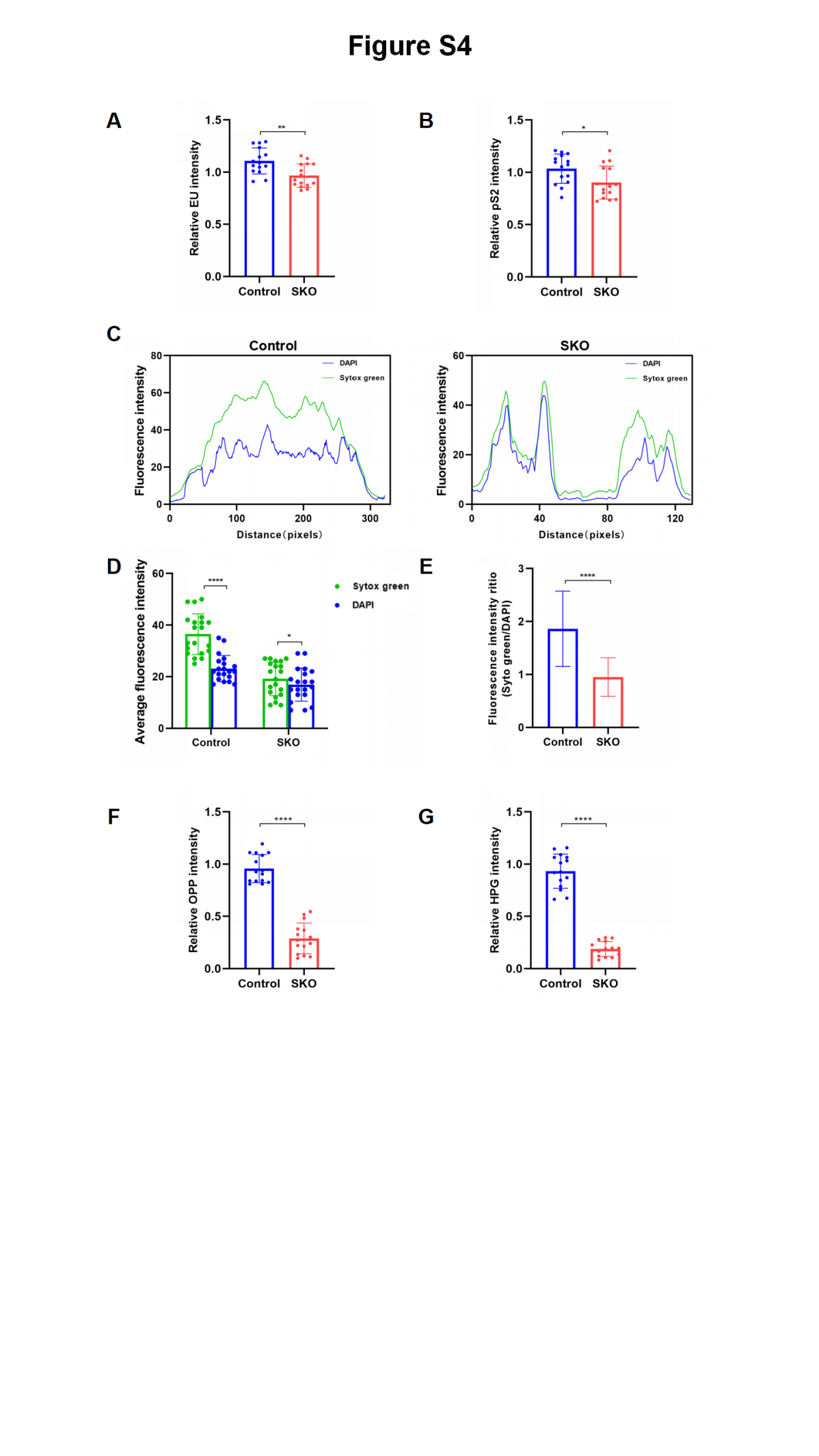

**Figure S4. Evaluation of transcription, translation, and nucleic acid distribution in 2-cell embryos**

(A) Quantification of EU signal intensity in late 2-cell embryos of *Cwf19l2*-SKO and control mice. Data are presented as the mean ± SD, n=15, ns: not significan.
(B) Quantification of pS2 signal intensity in late 2-cell embryos of *Cwf19l2*-SKO and control mice. Data are presented as the mean ± SD, n=15, ns: not significan.
(C) Intensity profiles of DAPI (blue) and Sytox green (green) in late 2-cell embryos ( Figure 4F) of *Cwf19l2*-SKO and control mice .
(D) Quantification of DAPI and Sytox green signal intensity in late 2-cell embryos of *Cwf19l2*-SKO and control mice. Data are presented as the mean ± SD. n=20, ****:*P* < 0.0001, ns: not significant.
(E) Ritio of DAPI and Sytox green signal intensity in late 2-cell embryos of *Cwf19l2*-SKO and control mice. Data are presented as the mean ± SD. ****:*P* < 0.0001.
(F) Quantification of OPP signal intensity in late 2-cell embryos of *Cwf19l2*-SKO and control mice. Data are presented as the mean ± SD, n=15, ****:*P* < 0.0001.
(G) Quantification of HPG signal intensity in late 2-cell embryos of *Cwf19l2*-SKO and control mice. Data are presented as the mean ± SD, n=15, ****:*P* < 0.000

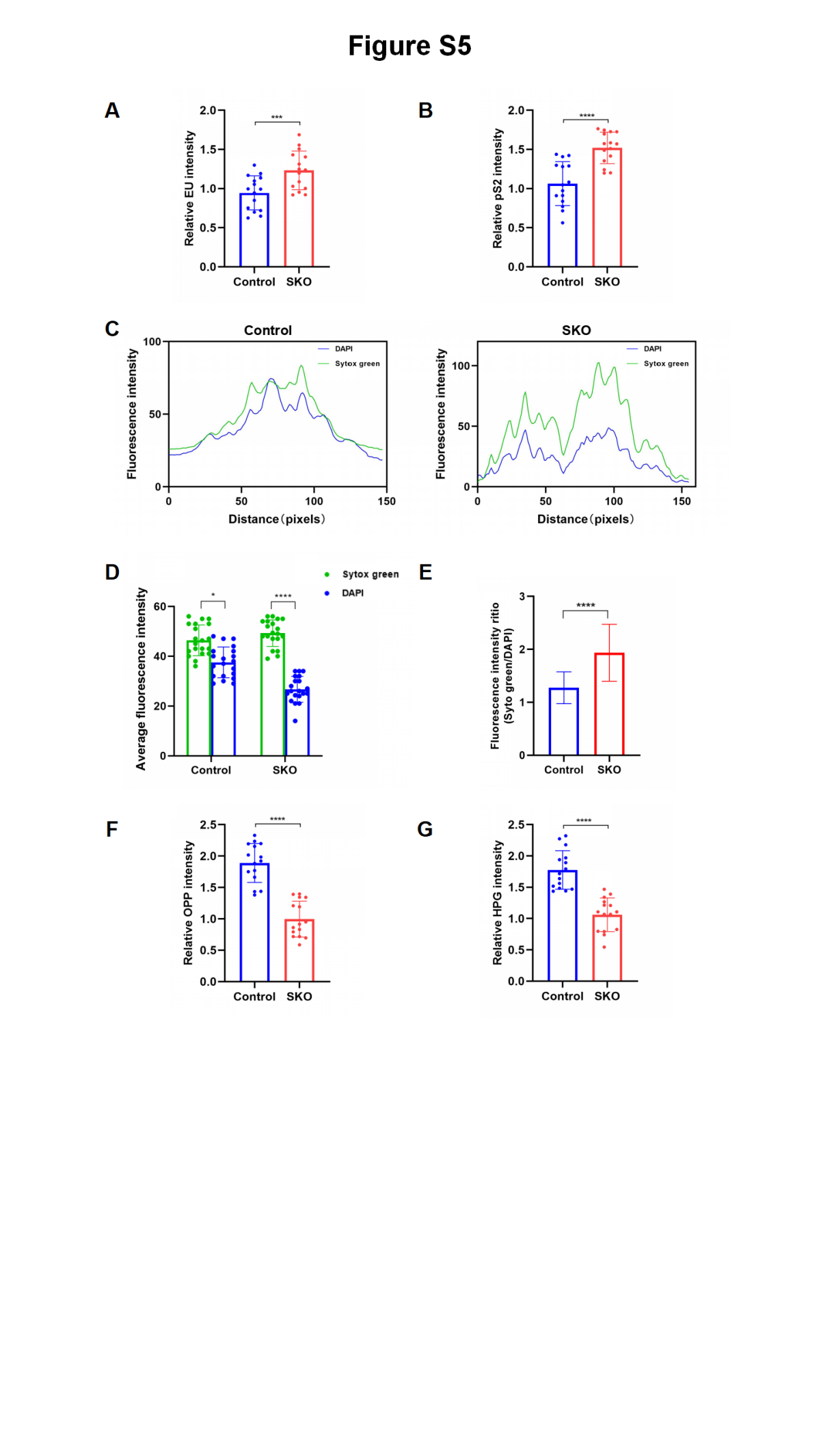

**Figure S5. Evaluation of transcription, translation, and nucleic acid distribution in GV oocytes**

(A) Quantification of EU signal intensity in GV oocytes of *Cwf19l2*-SKO and control mice. Data are presented as the mean ± SD, n=15, ns: not significan.
(B) Quantification of pS2 signal intensity in GV oocytes of *Cwf19l2*-SKO and control mice. Data are presented as the mean ± SD, n=15, ns: not significan.
(C) Intensity profiles of DAPI (blue) and Sytox green (green) in GV oocytes (Figure 4L) of *Cwf19l2*-SKO and control mice.
(D) Quantification of DAPI and Sytox green signal intensity in GV oocytes of *Cwf19l2*-SKO and control mice. Data are presented as the mean ± SD. n=20, *:*P* < 0.05, ****:*P* < 0.0001.
(E) Ritio of DAPI and Sytox green signal intensity in GV oocytes of *Cwf19l2*-SKO and control mice. Data are presented as the mean ± SD. ****:*P* < 0.0001.
(F) Quantification of OPP signal intensity in GV oocytes of *Cwf19l2*-SKO and control mice. Data are presented as the mean ± SD, n=15, ****:*P* < 0.0001.
(G) Quantification of HPG signal intensity in GV oocytes of *Cwf19l2*-SKO and control mice. Data are presented as the mean ± SD, n=15, ****:*P* < 0.0001.

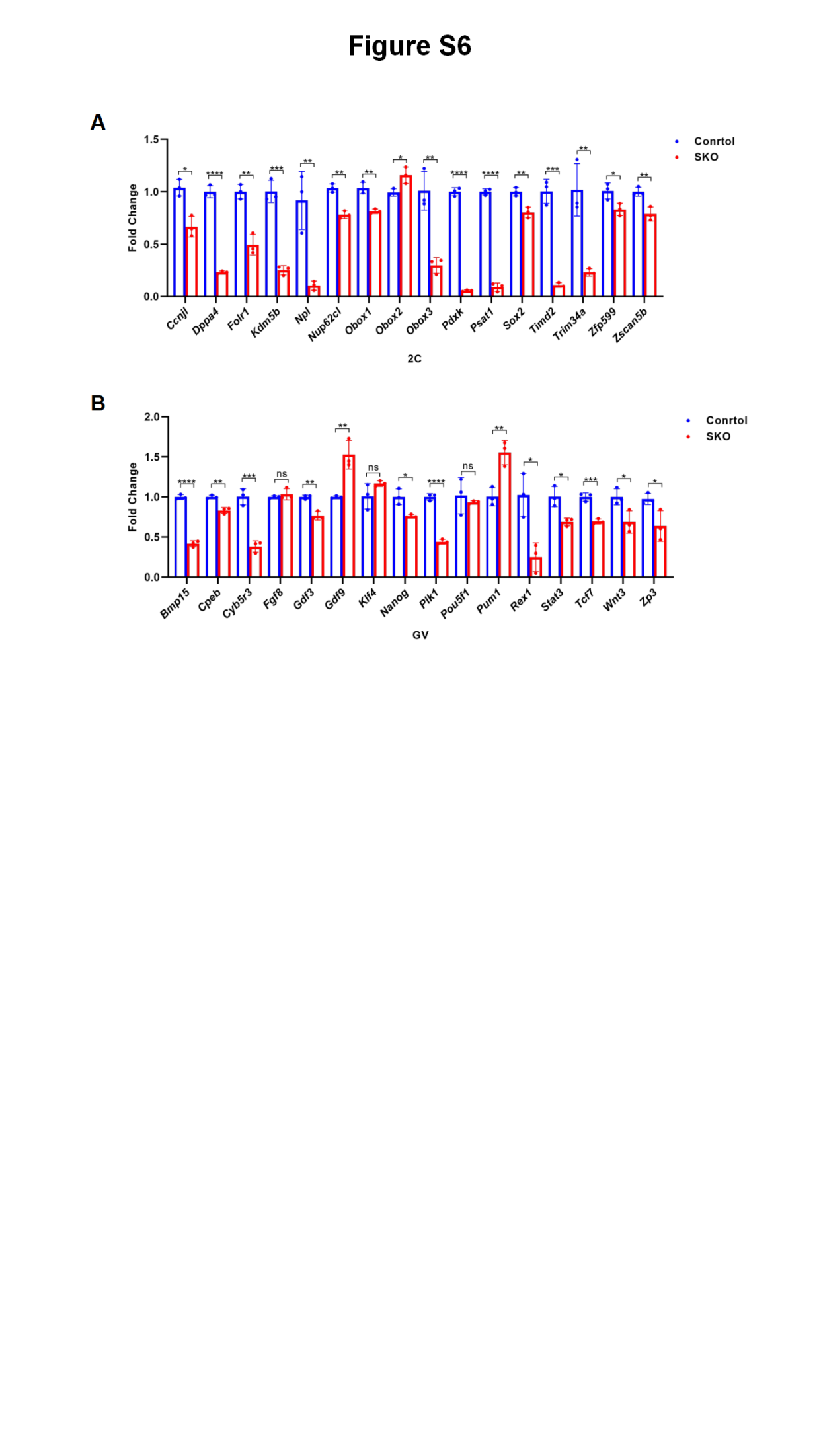

**Figure S6. Expression of key genes in GV oocytes and 2-cell embryos**

(A) QPCR analysis of maternal factors-related genes in GV oocytes of *Cwf19l2*-SKO and control mice. Data are presented as mean ± SD, n = 3, *P < 0.05, **P < 0.01, ***P < 0.001, ****P < 0.0001.
(B) QPCR analysis of ZGA-related genes in late 2-cell embryos of *Cwf19l2*-SKO and control mice. Data are presented as mean ± SD, n = 3, *P < 0.05, **P < 0.01, ***P < 0.001, ****P < 0.0001.

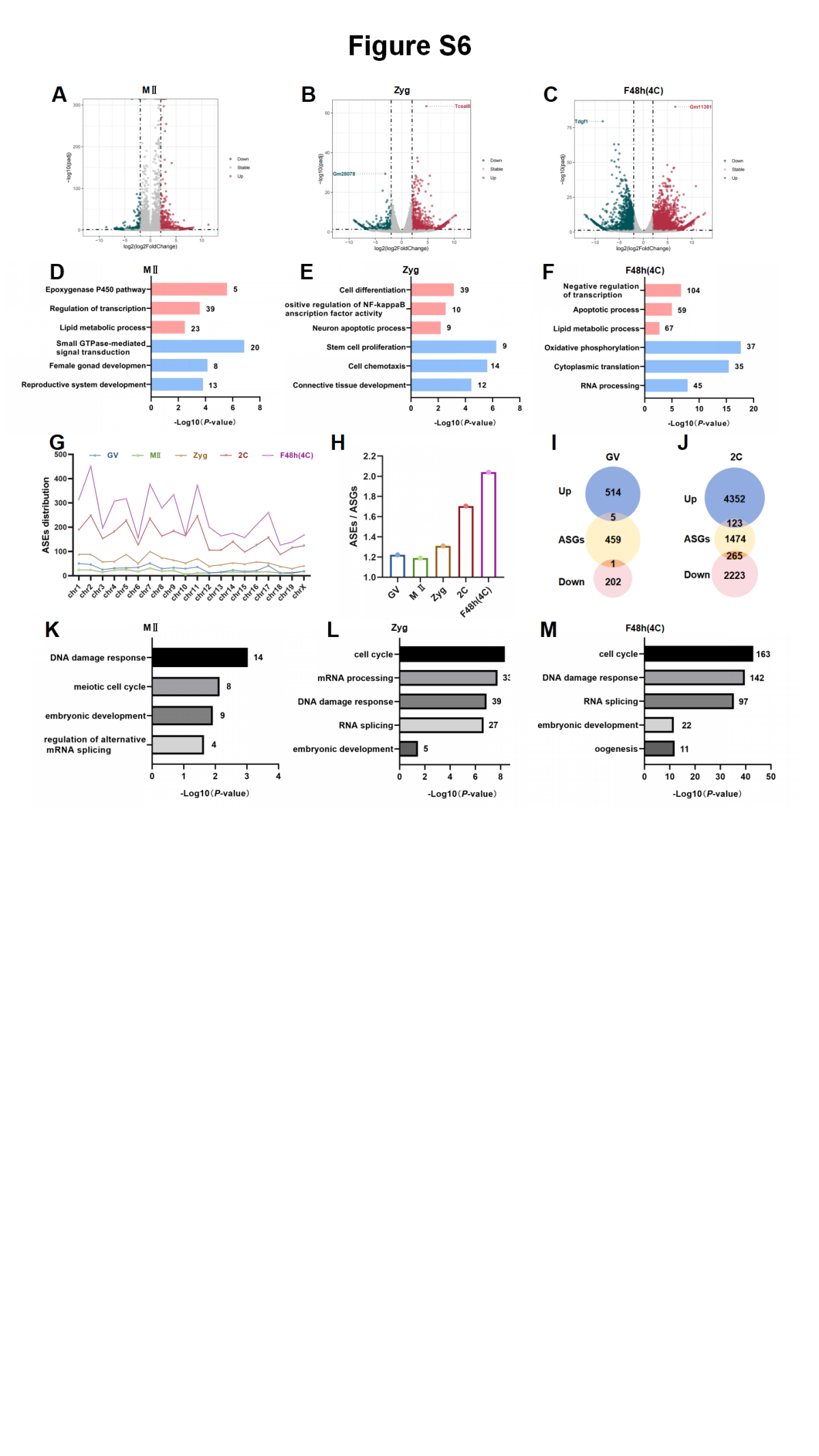

**Figure S7. Deficiency of CWF19L2 resulted in transcriptional disorganization across multiple developmental stages of oocytes and embryos**

(A) Volcano plot of DEGs identified by Smart-seq2 in MII oocytes of *Cwf19l2*-SKO and control mice.
(B) Volcano plot of DEGs identified by Smart-seq2 in zygotes of *Cwf19l2*-SKO and control mice.
(C) Volcano plot of DEGs identified by Smart-seq2 in 48 h post-fertilization embryos of *Cwf19l2*-SKO and control mice.
(D) GO analysis of DEGs in MII oocytes.
(E) GO analysis of DEGs in zygotes.
(F) GO analysis of DEGs in 48 h post-fertilization embryos.
(G) Chromosomal distribution of aberrant ASEs across 5-stage oocytes and early embryos.
(H) Ratio of aberrant ASEs to ASGs across 5-stage oocytes and early embryos.
(I) Venn diagram showing the common genes between DEGs and ASEs in GV oocytes.
(J) Venn diagram showing the common genes between DEGs and ASEs in 2-cell embryos.
(K) GO analysis of ASEs in MII oocytes.
(L) GO analysis of ASEs in zygotes.
(M) GO analysis of ASEs in 48 h post-fertilization embryos.

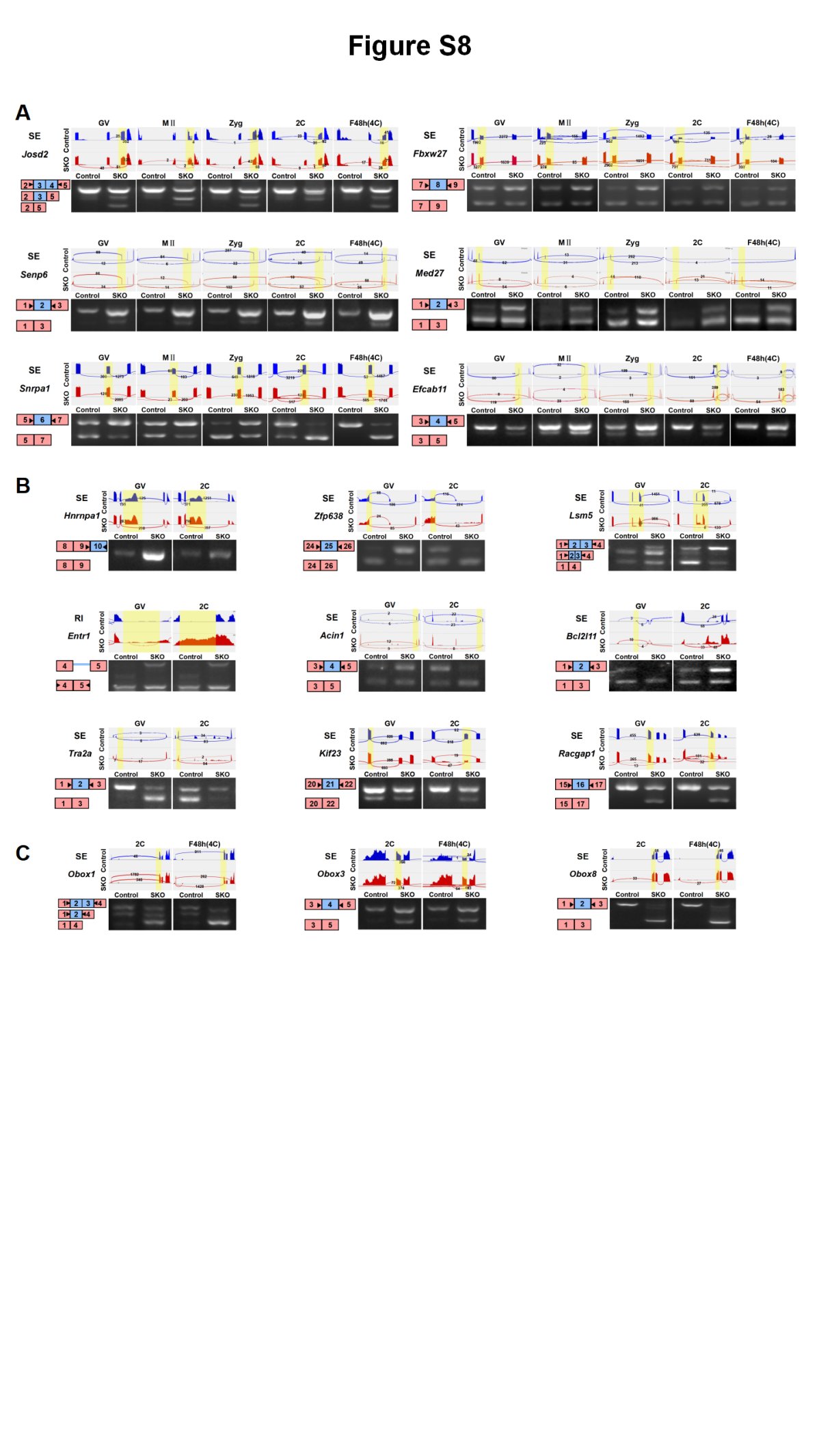

**Figure S8. Visualization and validation of abnormal ASEs in *Cwf19l2*-SKO and control mice**

(A) Visualization and validation of abnormal ASEs identified by Smart-seq2 in 5-stage oocytes and early embryos from *Cwf19l2*-SKO and control mice.
(B) Visualization and validation of abnormal ASEs identified by Smart-seq2 in GV oocytes and 2-cell embryos from *Cwf19l2*-SKO and control mice.
(C) Visualization and validation of abnormal ASEs identified by Smart-seq2 in 2-cell and 48 h post-fertilization embryos from *Cwf19l2*-SKO and control mice.

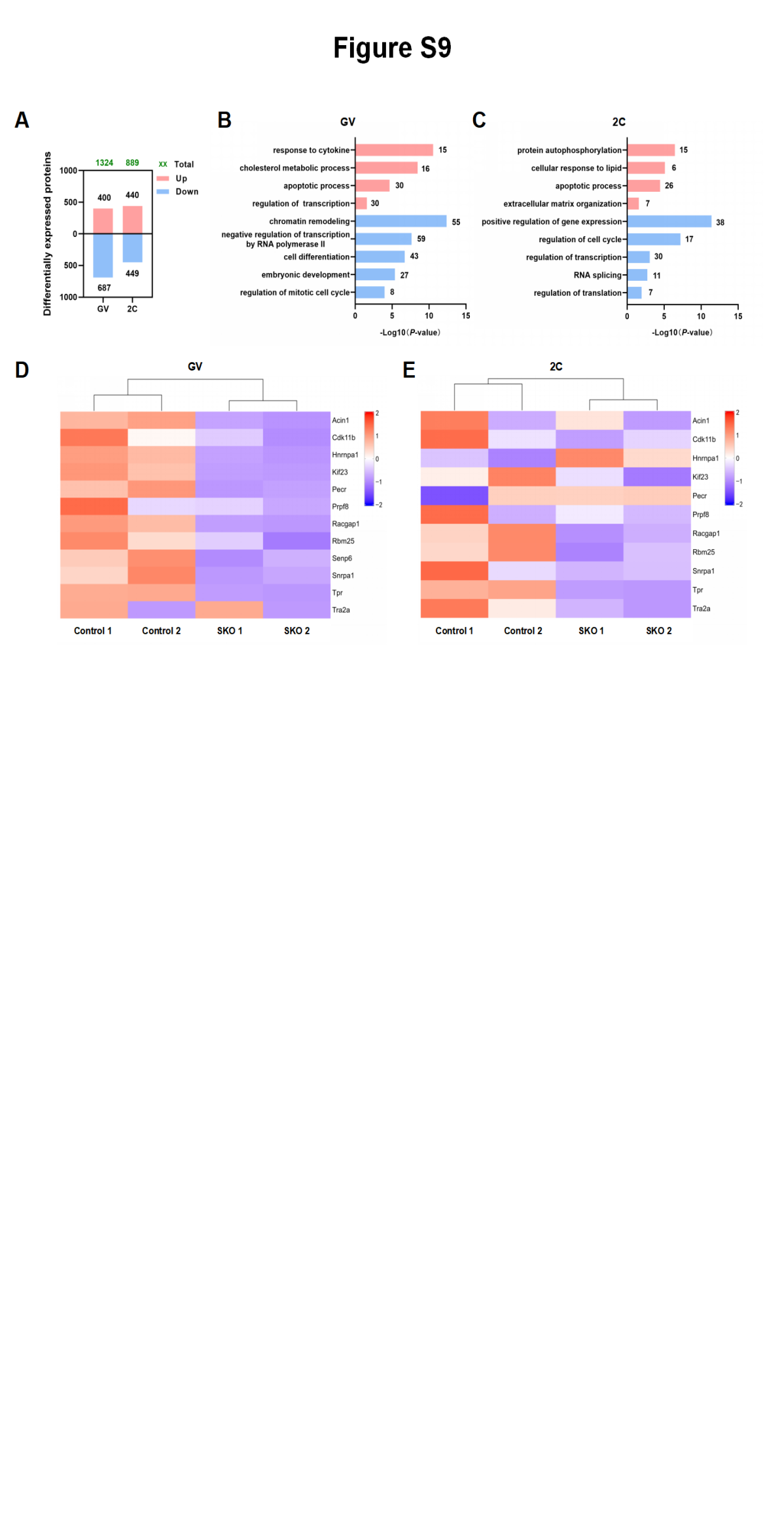

**Figure S9. Proteomic results of GV oocytes and 2-cell embryos from *Cwf19l2*-SKO and control mice**

(A) Number of DEPs in GV oocytes and 2-cell embryos. Red indicates upregulated proteins; blue indicates downregulated proteins; and green indicates total number.
(B) GO analysis of DEPs in GV oocytes. Red indicates upregulated proteins; blue indicates downregulated proteins.
(C) GO analysis of DEPs in 2-cell embryos.
(D) Heatmap of protein expression for genes with ASEs in GV oocytes.
(E) Heatmap of protein expression for genes with ASEs in 2-cell embryos.

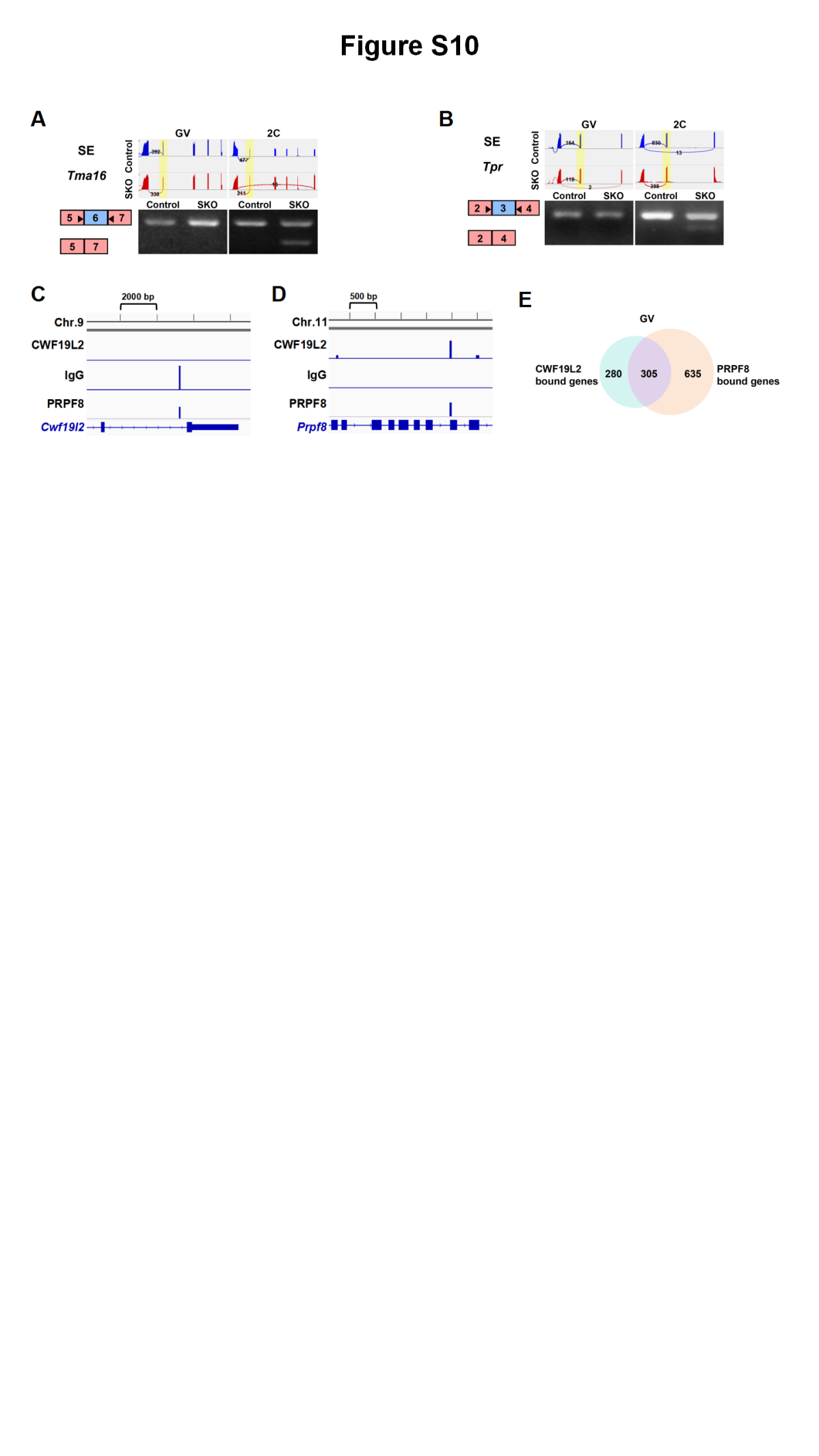

**Figure S10.CWF19L2 directly regulates alternative splicing and indirectly controls global gene splicing programs via modulation of additional splicing factors**

(A) Visualization and validation of *Tma16* abnormal ASEs in GV oocytes and 2-cell embryos from *Cwf19l2*-SKO and control mice.
(B) Visualization and validation of *Tpr* abnormal ASEs in GV oocytes and 2-cell embryos from *Cwf19l2*-SKO and control mice.
(C) Genome browser tracks showing LACE-seq binding peak distributions of CWF19L2, PRPF8, and IgG in *Cwf19l2* locus in GV oocytes.
(D) Genome browser tracks showing LACE-seq binding peak distributions of CWF19L2, PRPF8, and IgG at the *Prpf8* locus in GV oocytes.
(E) Venn diagram showing the common genes between CWF19L2 and PRPF8 bound genes in GV oocytes.

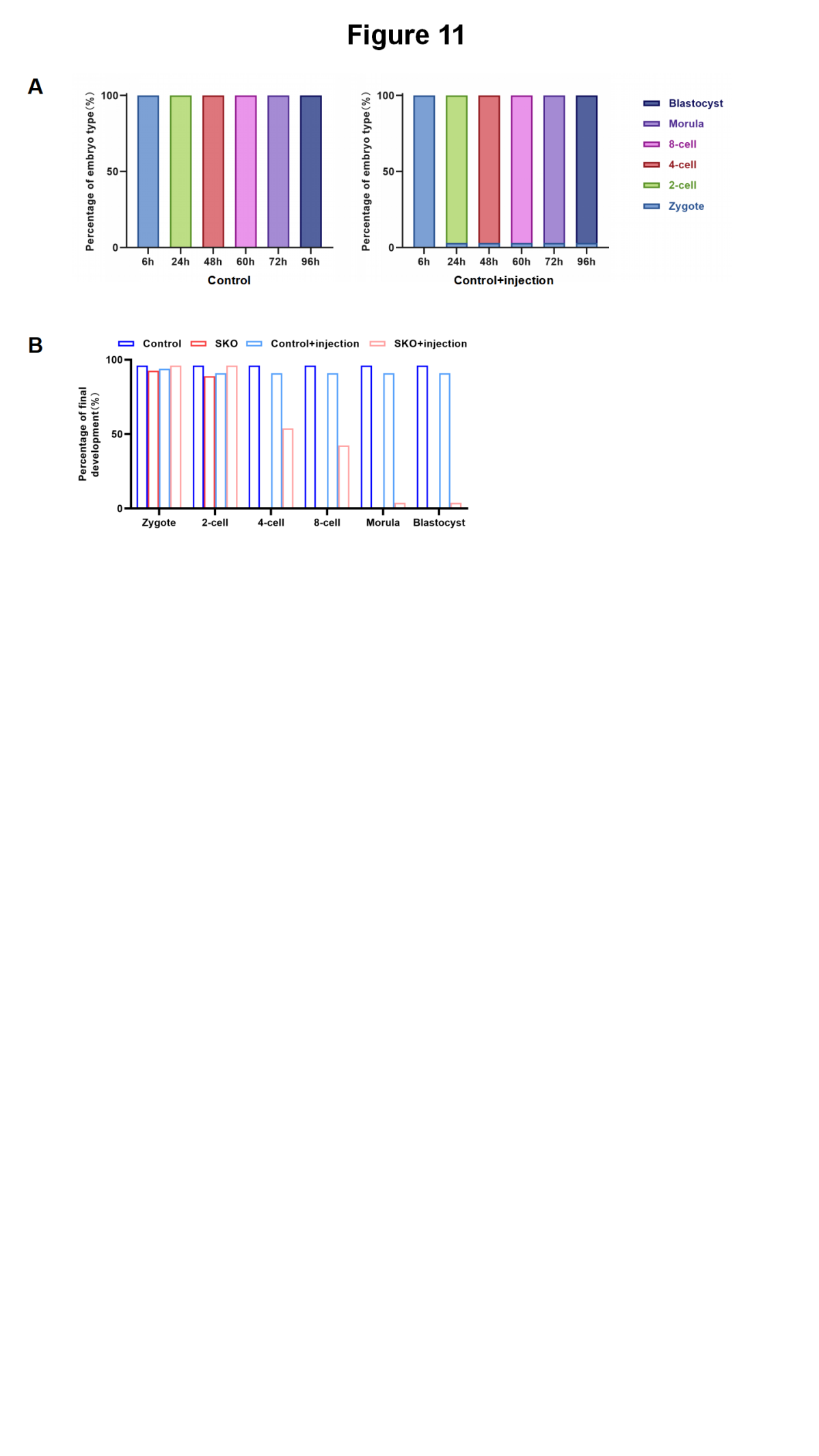

**Figure S11. Functional consequences of *Cwf19l2* restoration in CWF19L2-deficient embryos**

(A) The percentage composition of different embryo types in adult control mice with or without injection at 6, 24 48, 60, 72 and 96h after fertilization.
(B) Quantification of pre-implantation embryos at different stages derived from *Cwf19l2*-SKO and control mice without or with injection at 96h after fertilization.
